# Local RNA translation controls cell migration and Rab GTPase function

**DOI:** 10.1101/2020.05.19.104463

**Authors:** Konstadinos Moissoglu, Michael Stueland, Alexander N. Gasparski, Tianhong Wang, Lisa M. Jenkins, Michelle L. Hastings, Stavroula Mili

## Abstract

Numerous RNAs exhibit specific distribution patterns in mammalian cells. However, the functional and mechanistic consequences are relatively unknown. We investigate here the functional role of RNA localization at cellular protrusions of mesenchymal migrating cells, using as a model the *RAB13* RNA, which encodes a GTPase important for vesicle-mediated membrane trafficking. While *RAB13* RNA is enriched at peripheral protrusions, the expressed protein is concentrated perinuclearly. By specifically preventing *RAB13* RNA localization, we show that peripheral *RAB13* translation is not important for the overall distribution of the RAB13 protein, or its ability to associate with membranes, but is required for full activation of the GTPase and for efficient cell migration. This effect is mediated by a co-translational association of RAB13 with the exchange factor RABIF. Our results indicate that RAB13-RABIF association at the periphery is required for directing RAB13 GTPase activity to promote cell migration. Thus, translation of RAB13 in specific subcellular environments imparts the protein with distinct properties and highlights a means of controlling protein function through local RNA translation.

## INTRODUCTION

Localization of RNAs to diverse subcellular destinations is widely observed in various cell types and organisms (Buxbaum et al., 2015, Medioni et al., 2012, Meignin and Davis, 2010). However, in mammalian cells, the functional and mechanistic consequences are relatively unknown.

In some cases, RNA accumulation can be accompanied by a corresponding increase in protein concentration at the same location. Such local protein gradients can be reinforced through translationally silencing RNAs prior to arrival at their destination (Besse and Ephrussi, 2008), thus ensuring tight spatial and temporal control of protein production and preventing deleterious effects of premature or ectopic translation (Buxbaum et al., 2015, Jung et al., 2014). This type of regulation has been described in highly polarized cells, such as neurons. For example, translational activation of RNAs localized at growth cones, and the consequent increase in local protein abundance, underlie axonal pathfinding decisions (Colak et al., 2013, Leung et al., 2006b, Wong et al., 2017). Similarly, activation of dendritic synapses upregulates translation of localized transcripts and is important for synaptic plasticity (Holt et al., 2019, Rangaraju et al., 2017, Yoon et al., 2016). Indeed, RNA localization appears to direct enrichment to neurites of almost half of the neurite-enriched proteome (Zappulo et al., 2017). A similar significant correlation between steady-state RNA and protein localization has been described in epithelial cells for proteins associated with organelles, such as mitochondria and the endoplasmic reticulum (Fazal et al., 2019).

Nevertheless, a concordance between RNA localization and protein distribution is not always observed. One case in point concerns RNAs enriched at dynamic protrusions of mesenchymal migrating cells. RNA localization at protrusions is important for protrusion stability and cell migration (Mardakheh et al., 2015, Mili et al., 2008, Wang et al., 2017). However there is little correlation between RNA and protein distributions (Mardakheh et al., 2015) and protrusion-enriched RNAs can be similarly translated in both internal and peripheral locations (Moissoglu et al., 2019), thus raising the question of what the functional role of RNA transport in these cases is. Here we investigate the consequences of local peripheral translation focusing on the *RAB13* RNA.

RAB13 is a member of the Rab family of small GTPases which play important roles in vesicle-mediated membrane trafficking (Ioannou and McPherson, 2016, Pfeffer, 2017). It is amplified in the majority of cancers and its levels inversely correlate with prognosis (Ioannou and McPherson, 2016). Activation of RAB13 at the plasma membrane is required for cell migration and invasion (Ioannou et al., 2015), potentially through multiple mechanisms, including activity-dependent recycling of integrins or modulation of actin-binding proteins at the leading edge (Sahgal et al., 2019, Sakane et al., 2012, Sakane et al., 2013). *RAB13* RNA is prominently localized at protrusive regions of multiple cell types (Feltrin et al., 2012, Mili et al., 2008, Moissoglu et al., 2019) together with a group of RNAs whose localization is regulated by the Adenomatous Polyposis Coli (APC) protein and detyrosinated microtubules (Wang et al., 2017)

We show here that *RAB13* RNA and protein distributions are quite discordant, with *RAB13* RNA being enriched in the periphery while RAB13 protein assumes mostly a perinuclear distribution. To assess the functional role of peripheral RNA localization, we devise a way to specifically prevent localization of *RAB13* RNA at peripheral protrusions without affecting its translation, stability or the localization of other co-regulated RNAs. Importantly, we show that peripheral RAB13 translation does not affect the overall distribution of the protein or its ability to associate with membranes but is required for activation of the GTPase and for efficient cell migration. Our data show that RAB13 associates co-translationally with the exchange factor RABIF. Peripheral translation is required for RABIF-RAB13 interaction at the periphery and for directing RAB13 GTPase activity to promote cell migration. Our results indicate that translation of RAB13 in specific subcellular environments imparts the protein with distinct properties, thus highlighting a means of controlling protein function through local RNA translation.

## RESULTS

### RAB13 RNA and protein exhibit distinct subcellular distributions

In both mouse and human mesenchymal cells, *RAB13* RNA is prominently enriched at peripheral protrusions (Figure 1A and (Mili et al., 2008, Wang et al., 2017)), where it can be actively translated (Moissoglu et al., 2019). To assess whether this leads to a corresponding increase of RAB13 protein, we visualized the distribution of endogenous RAB13. Interestingly, despite the peripheral *RAB13* RNA enrichment, RAB13 protein is prominently concentrated around the nucleus (Figure 1B, C). To independently address this, we fractionated protrusions (Ps) and cell bodies (CB) from cells grown on microporous filters and assessed protein and RNA distributions between them (Figure 1D). Consistent with the imaging data above, *RAB13* RNA is significantly enriched at protrusions while RAB13 protein is not (Figure 1D). Taken together, these results suggest that at least a significant proportion of the protein translated from peripheral *RAB13* RNA does not persist at the periphery but assumes a steady state perinuclear distribution.

**Figure 1:**
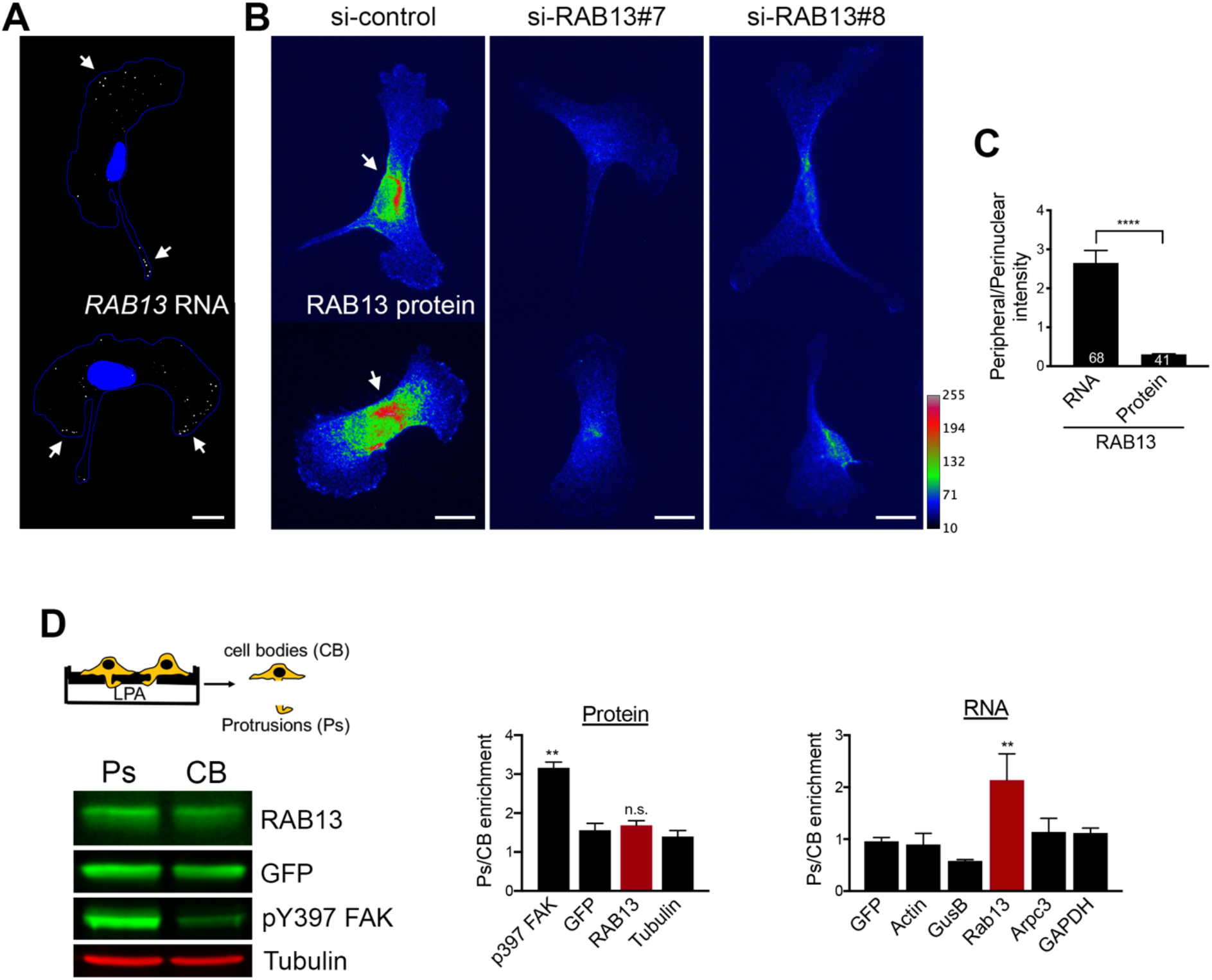
RAB13 RNA and protein exhibit distinct subcellular distributions. **A.** Representative FISH images showing *RAB13* RNA distribution in MDA-MB-231 cells. Nuclei and cell outlines are shown in blue. Arrows points to *RAB13* RNA concentrated at protrusive regions. **B.** Representative immunofluorescence images of RAB13 protein in cells transfected with the indicated siRNAs. Reduction of intensity in RAB13 knockdown cells confirms the specificity of the signal. Arrows point to perinuclear RAB13 protein. Calibration bar shows intensity values. **C.** Ratios of peripheral/perinuclear intensity calculated from images as shown in A and B. Values within each bar represent number of cells observed in 3 independent experiments. **D.** Protrusions (Ps) and cell bodies (CB) of cells induced to migrate towards LPA were isolated and analyzed to detect the indicated proteins (by Western blot; left panels) or RNAs (by RT-ddPCR; right panel). Ps/CB enrichment ratios from 2 independent experiments are shown. The enrichment of pY397-FAK serves to verify the enrichment of protrusions containing newly formed adhesions in the Ps fraction. P-values: **<0.01; ****<0.0001 by student’s t-test (C) or analysis of variance with Dunnett’s multiple comparisons test (D). Scale bars: 10 μm.

### A GA-rich motif within the mouse Rab13 3’UTR is necessary for localization at protrusions

To understand the functional role of peripheral translation, we first sought to narrow down on specific localization sequences. We had previously shown that a 200-300nt region of the mouse *Rab13* 3’UTR is sufficient for localization and can competitively inhibit the localization of other peripheral, APC-dependent RNAs (Wang et al., 2017), suggesting that it contains a binding site for a factor commonly bound to APC-dependent RNAs. Using sequence alignment and gazing we noticed a particular GA-rich motif, with the consensus RGAAGRR (where R is a purine), which is present, in one or multiple copies, in the 3’UTR of the majority (~60%) of APC-dependent RNAs (Figure 2A and S1) and which is significantly enriched (p=4.99e-5; motif enrichment analysis (meme-suite.org)) in APC-dependent RNAs compared to APC-independent RNAs, an RNA group which is also enriched at protrusions but through a distinct pathway (Wang et al., 2017). To test for any functional significance, we expressed an exogenous RNA carrying either the wild type Rab13 3’UTR, or the 3’UTR carrying specific deletions of this motif (Figure 2A, B). We imaged RNAs using single-molecule FISH and measured a Peripheral Distribution Index (PDI) to quantify their distributions in multiple cells (Stueland et al., 2019). Consistent with previous observations (Wang et al., 2017), a control β-globin RNA shows a mostly diffuse cytoplasmic distribution, (Figure 2B, C), while addition of the Rab13 3’UTR is sufficient to promote its peripheral localization, denoted by low and high PDI values, respectively. Interestingly, deletion of one RGAAGRR motif (Rab13 UTR (Δ1)) significantly perturbed the ability of the Rab13 3’UTR to direct localization of the β-globin RNA, while deletion of two of them (Rab13 UTR (Δ1+2)) made the distribution of the reporter indistinguishable from that of the non-localized control (Figure 2C). An endogenous localized RNA (*Ddr2*) remained similarly localized at protrusions in all conditions (Figure 2C). Therefore, at least some of the RGAAGRR motifs are required for peripheral localization.

**Figure 2:**
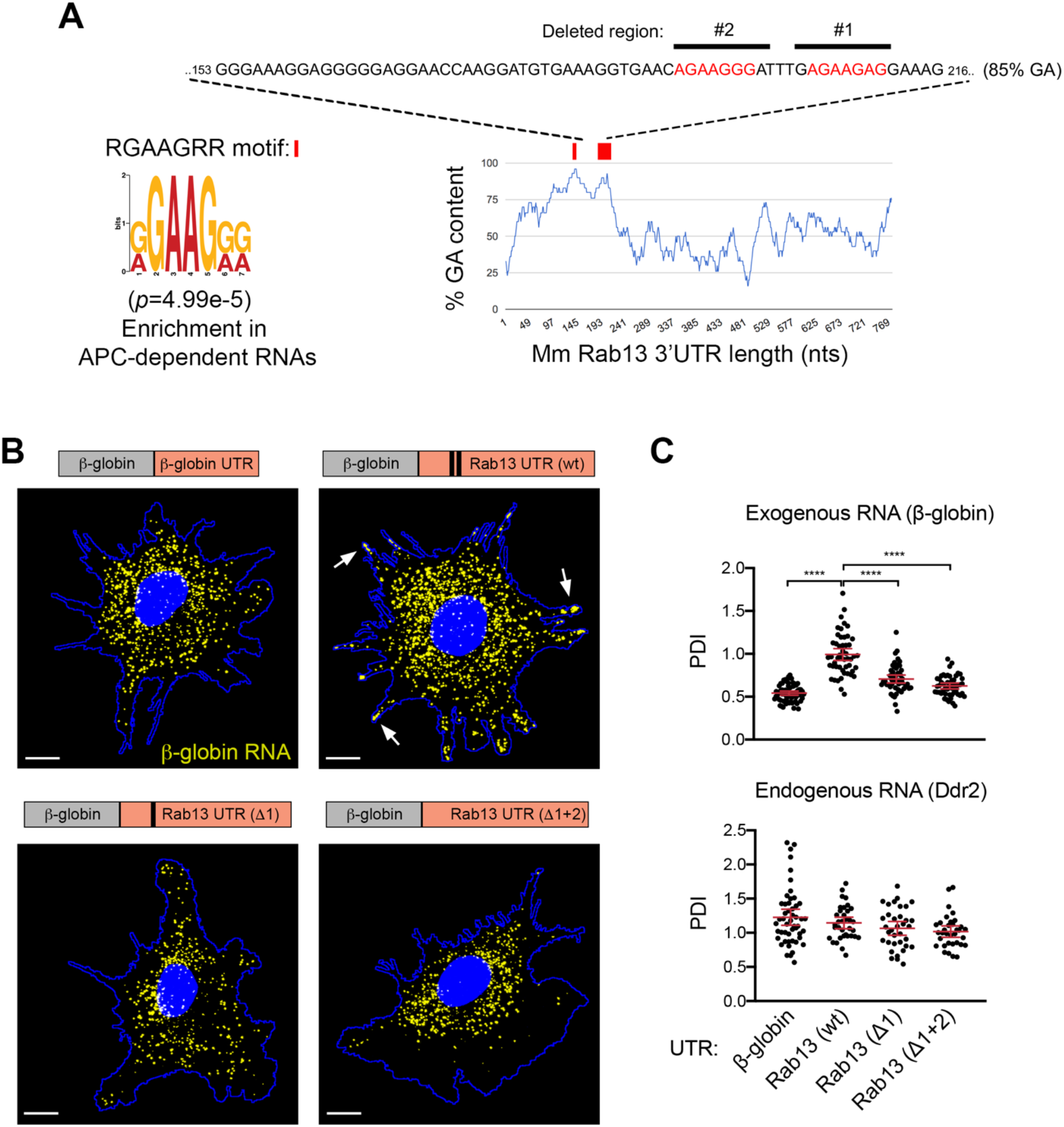
A GA-rich motif in the mouse *Rab13* 3’UTR is necessary for localization at protrusions. **A.** Schematic showing the %GA content along the mouse *Rab13* 3’UTR using a 30nt window size. Occurrences of the consensus GA-rich motif are indicated by a red rectangle. The exact sequence between nucleotides 153-216 is shown with GA motifs in red and deleted regions indicated by black bars. **B.** FISH images of mouse fibroblasts expressing the β-globin coding sequence followed by the indicated UTRs. β-globin RNA is shown in yellow. Nuclei and cell outlines are shown in blue. Arrows points to β-globin RNA concentrated at protrusive regions. Δ1 and Δ1+2 indicate deletions of the regions shown in (A). Scale bars: 10 μm. **C.** Distribution of β-globin RNA, or of *Ddr2* RNA detected in the same cells, quantified by measuring a Peripheral Distribution Index (PDI). N=35-55 cells observed in 3 independent experiments. ****:p<0.0001 by analysis of variance with Dunnett’s multiple comparisons test.

### Antisense oligonucleotides against the GA-rich region specifically interfere with localization of Rab13 RNA

Another notable feature of these motifs is that the majority of them (62%; 148 of 239 motifs in 3’UTRs of mouse APC-dependent RNAs) are found within more extended GA-rich regions, which exhibit high GA content (>75%) for at least 30 consecutive nucleotides or more (Figure S1). To further investigate the roles of these different features and to, at the same time, interfere with the localization of the endogenous *Rab13* RNA, we used antisense oligonucleotides (ASOs), which can interfere with RNA structure formation or RNA-protein binding (Havens and Hastings, 2016, Hua et al., 2010, Lentz et al., 2013) (Figure 3A). Here, we utilized 25nt-long phosphorodiamidate morpholino (PMO) ASOs.

**Figure 3:**
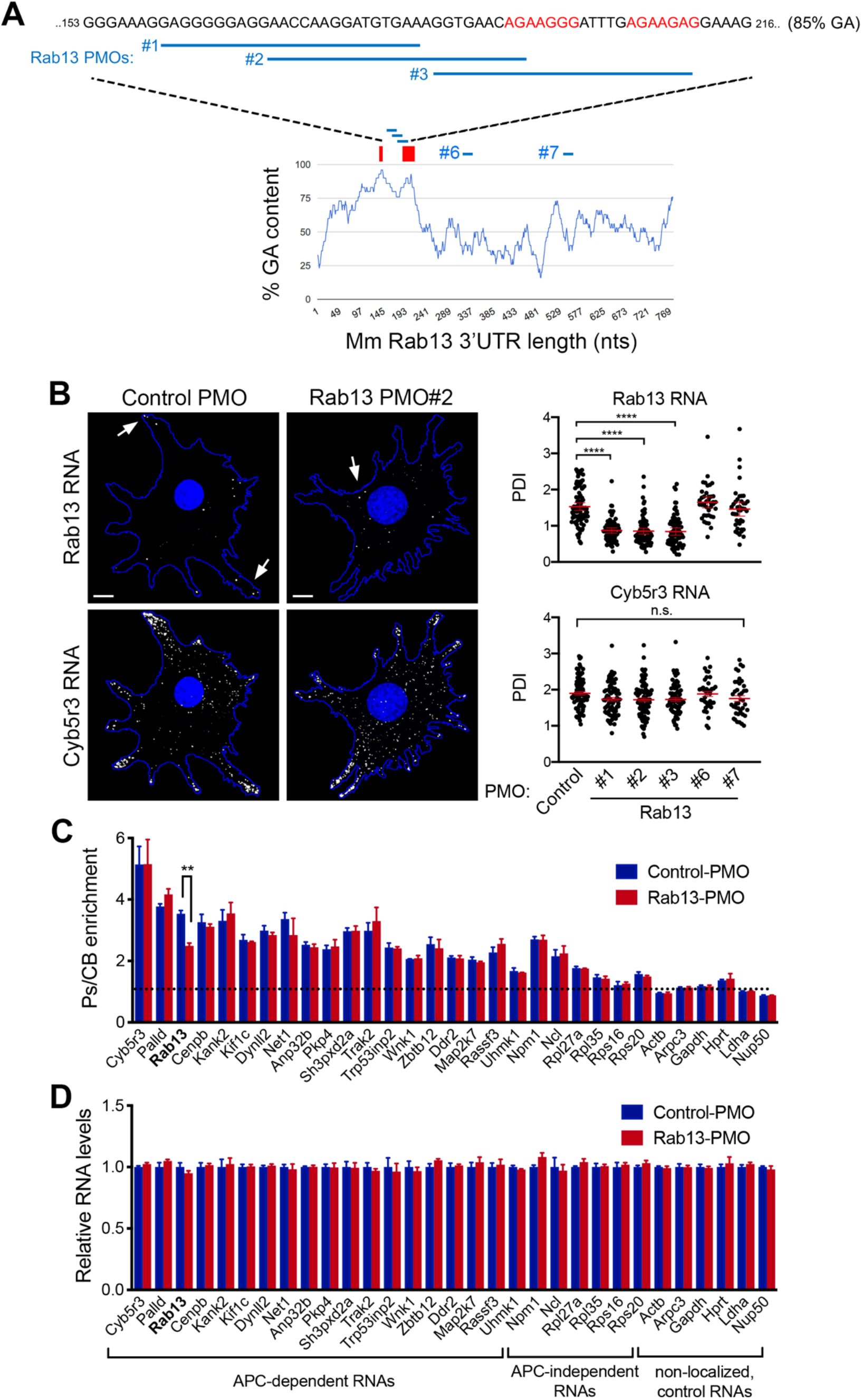
Antisense oligonucleotides against the GA-rich region specifically interfere with localization of *Rab13* RNA. **A.** Schematic showing positions along the mouse *Rab13* 3’UTR targeted by the indicated PMOs. PMOs #1, 2 and 3 target the RGAAGRR motifs or the adjacent GA-rich region. PMOs #6 and #7 target the *Rab13* 3’UTR outside of the GA-rich region. The control PMO targets an intronic sequence of human β-globin. Red rectangles and text indicate the location of the GA-rich motifs. **B.** FISH images and corresponding PDI measurements of mouse fibroblast cells treated with the indicated PMOs. *Cyb5r3* is an APC-dependent RNA also enriched at protrusions. Arrows point to locations of *Rab13* RNA accumulation. Note that *Rab13* RNA becomes perinuclear in cells treated with PMOs against the GA-rich region. Scale bars: 10 μm. ****: p<0.0001 by analysis of variance with Dunnett’s multiple comparisons test. N=40-90 cells observed in 3-6 independent experiments. **C.** Protrusion (Ps) and cell body (CB) fractions were isolated from cells treated with control-PMO or Rab13-PMO #2. The indicated RNAs were detected through nanoString analysis to calculate Ps/CB enrichment ratios (n=3; error bars: standard error). Note that only the distribution of *Rab13* RNA is affected. **: p=0.01 by two-way ANOVA with Bonferroni’s multiple comparisons test against the corresponding control. **D.** Levels of the indicated RNAs were determined using nanoString analysis from control- or Rab13 PMO #2-treated cells. N=4. No significant differences were detected by two-way ANOVA against the corresponding controls.

We first delivered fluorescently labeled PMOs to determine the efficiency of delivery and their persistence in cells. PMOs were taken up by virtually all cells and persisted, either within apparent endosomal structures or released into the cytosol, for more than 3 days (Figure S2A, B). The effect of antisense PMOs on the localization of *Rab13* RNA was assessed, 3 days after PMO delivery, by single-molecule FISH of the endogenous *Rab13* RNA and PDI calculation. As expected, cells exposed to the control PMO exhibited peripheral localization of *Rab13* RNA. Similarly, the Rab13 #6 and #7 PMOs did not affect *Rab13* RNA distribution (Figure 3B). However, PMOs targeting the RGAAGRR motifs (PMOs #2 and #3) caused a pronounced mislocalization of *Rab13* RNA towards the perinuclear cytoplasm, evidenced by a significant reduction in PDI values (Figure 3B). Interestingly, the Rab13 PMO #1, which targets the adjacent GA-rich region, disrupted localization to a similar extent, suggesting that apart from the RGAAGRR motifs additional GA-rich sequences are important for localization or that the overall structure of this region is important. Notably, within the same cells, another APC-dependent RNA, *Cyb5r3*, maintained its localization at protrusions under all conditions. Therefore, PMOs against the GA-rich region of *Rab13* RNA appear to specifically perturb *Rab13* RNA localization at protrusions.

To more extensively investigate the specificity of the observed effect, we assessed the distribution of a panel of ~20 APC-dependent RNAs, as well as of several APC-independent RNAs, using a protrusion/cell body fractionation scheme (Wang et al., 2017). As described previously (Wang et al., 2017), APC-dependent RNAs are enriched at protrusions, and their enrichment is more pronounced than that exhibited by APC-independent RNAs (Figure 3C). Importantly, cells treated with a mislocalizing Rab13 PMO exhibited indistinguishable distributions for all RNAs tested, with the notable exception of the targeted *Rab13* RNA, which became significantly less enriched at protrusions, corroborating and extending the FISH analysis described above (Figure 3B). We conclude that antisense PMOs against the Rab13 GA-rich region, specifically alter the distribution of *Rab13* RNA without impacting the distribution of other RNAs, even those belonging to the same co-regulated group.

We additionally examined the overall abundance of the same panel of RNAs. PMO oligos do not trigger RNase H activity, and consistent with that, we did not observe any detectable change in the total levels of either *Rab13*, or of any other RNA, in cells treated with Rab13 PMOs (Figure 3D and S3). Therefore, this approach allows us to specifically alter the distribution of the endogenous *Rab13* RNA without affecting its overall abundance in cells.

### The human RAB13 3’UTR exhibits a functionally conserved GA-rich region required for peripheral localization

*RAB13* RNA is localized at protrusive regions in diverse cell types and species. We thus sought whether similar sequence determinants support localization of the human *RAB13* transcript, which exhibits a GA-rich region (>75%) and interspersed RGAAGRR motifs with similar topology as that of the mouse *Rab13* 3’ UTR sequence (nts 98-268) (Figure 4A). To identify functional regions with regards to RNA localization at protrusions, we delivered PMOs targeting regions across the length of the 3’UTR. PMOs targeting either RGAAGRR motifs directly (RAB13 PMOs 165 and 230) or adjacent GA-rich regions (RAB13 PMOs 91, 113, 191 and 210) (Figure 4A) significantly affected *RAB13* RNA localization. By contrast, all PMOs targeting sites outside of the GA-rich region did not affect RNA distribution (Figure 4A). Concomitant delivery of two individual PMOs (RAB13 PMOs 191 and 230) had an additive effect resulting in marked *RAB13* RNA mislocalization. Furthermore, the observed effects were specific for *RAB13* RNA since the distribution of another peripherally localized RNA, *NET1*, was not affected (Figure 4A, B). Thus, also in human cells, interfering with either the RGAAGRR motifs or the adjacent GA-rich regions specifically perturbs the peripheral localization of *RAB13* RNA.

**Figure 4:**
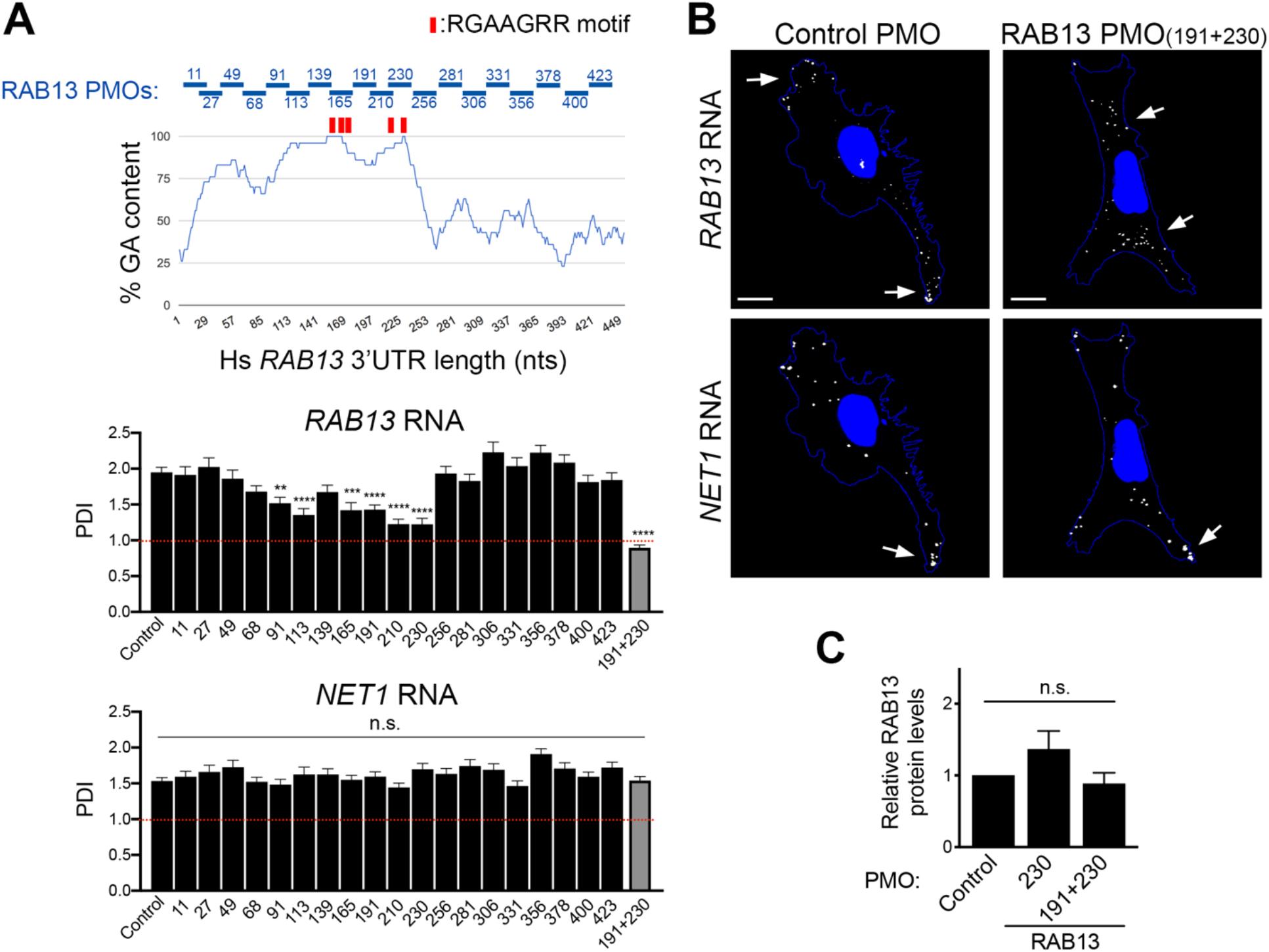
The human *RAB13* 3’UTR contains a functionally conserved GA-rich region required for peripheral localization. **A.** Schematic showing %GA content and positions along the human *RAB13* 3’UTR targeted by the indicated PMOs. Red rectangles indicate the location of GA-rich motifs. Graphs present PDI measurements of *RAB13* RNA (upper panel) or *NET1* RNA (another APC-dependent RNA; bottom panel) detected in MDA-MB-231 cells treated with the indicated PMOs. PDI=1 indicates a diffuse distribution. p-values: **<0.01, ***<0.001, ****<0.0001 by analysis of variance with Dunnett’s multiple comparisons test. n.s.: non-significant. N=30-73 cells observed in 3-5 independent experiments. **B.** Representative FISH images of cells treated with the indicated PMOs. Arrows point to locations of RNA accumulation. Note that *RAB13* RNA becomes perinuclear in cells treated with PMOs against the GA-rich region, while *NET1* remains localized at protrusions. Scale bars: 10 μm. **C.** RAB13 protein levels were measured by quantitative Western blot and normalized to total α-Tubulin or GAPDH levels. Relative levels in RAB13 PMO-treated compared to control are shown. No significant differences were detected by Kruskal-Wallis test with Dunn’s multiple comparisons test.

Importantly, RNA mislocalization was not accompanied by any detectable change in the amount of RAB13 protein produced (Figure 4C). Specifically, the same amount of RAB13 protein is produced in cells exhibiting either peripheral (control PMO) or perinuclear (RAB13 PMO 230 or 191+230) *RAB13* RNA distribution. This is consistent with the recently reported observation that *RAB13* RNA is similarly translated in both perinuclear and peripheral regions (Moissoglu et al., 2019). Therefore, the use of ASOs allows us to specifically assess the functional roles promoted by the localization of *RAB13* RNA without confounding contributions due to altered protein expression.

### Peripheral RAB13 RNA localization is important for cell migration

To understand the functional role of *RAB13* RNA localization at protrusions, we assessed the effect of *RAB13* RNA mislocalization on the ability of cells to migrate, given that the encoded RAB13 protein promotes cell migration through multiple mechanisms (Ioannou and McPherson, 2016). For this, we compared cells treated with control PMOs or RAB13 mislocalizing PMOs using various assays. In one case, cells plated on microporous Transwell membrane inserts were induced to migrate towards a chemoattractant gradient and the number of cells reaching the bottom surface after 4 hours was counted. Mislocalization of *RAB13* RNA from protrusions significantly perturbed the ability of cells to respond and migrate chemotactically (Figure 5A). To assess migration parameters of individual cells, single cells expressing Cherry-NLS fluorescent protein to mark nuclei were plated on collagen and were tracked over time to derive their speed and directionality (Figure 5B and movie S1). Again, cells containing perinuclear *RAB13* RNA exhibited lower migration speeds (Figure 5B). Finally, cells attached on one side of a Matrigel plug were induced to migrate through it towards a chemoattractant in order to assess their ability to invade through a 3-dimensional matrix. Serial imaging sections through the Matrigel were taken for >100 μm to assess the number of cells reaching at various depths. Notably, also in this case, cells treated with RAB13 mislocalizing PMOs exhibited significantly reduced invasiveness (Figure 5C). Therefore, peripheral localization of *RAB13* RNA functionally contributes to various cell migration modes.

**Figure 5:**
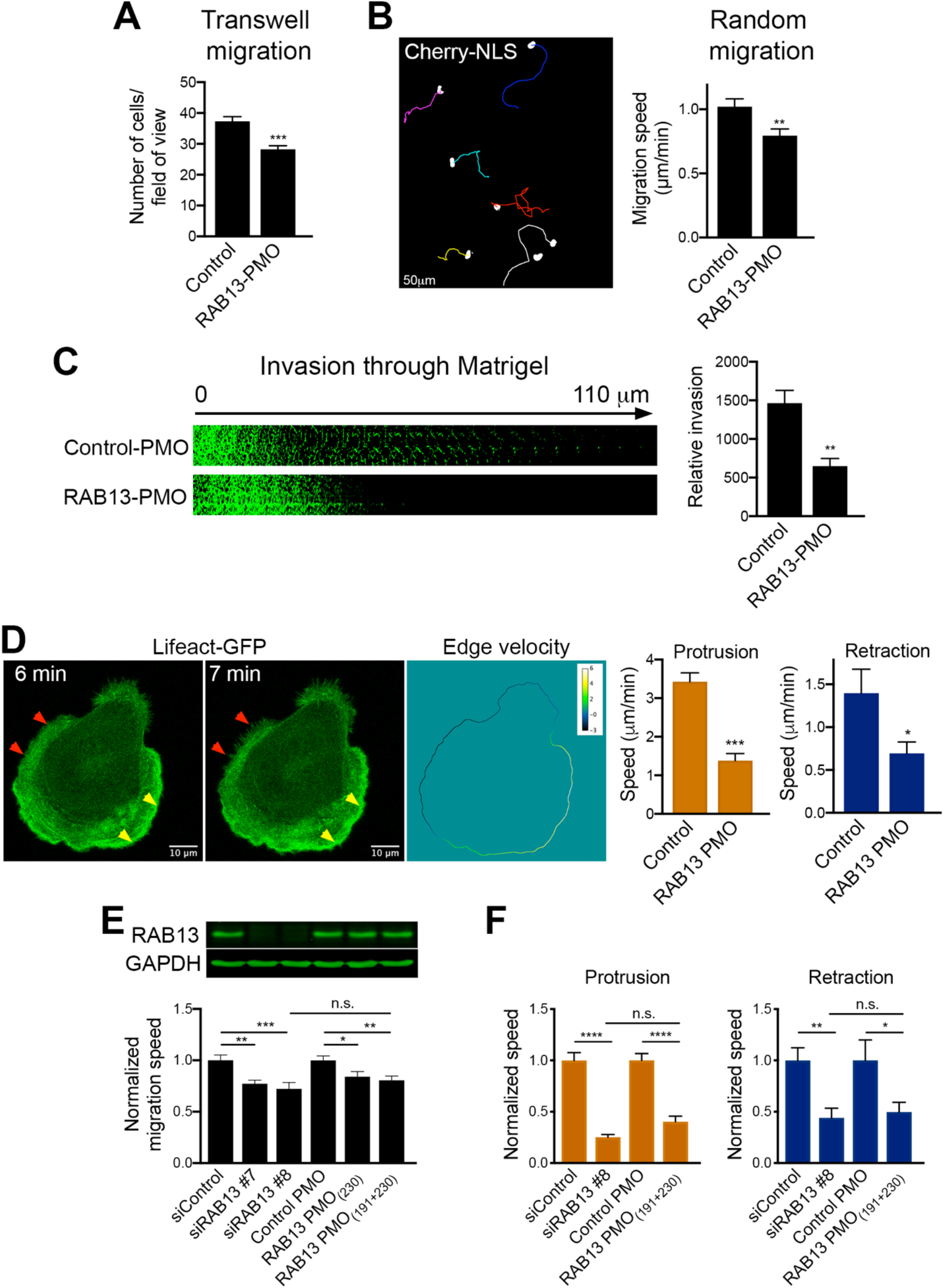
Loss of peripheral *RAB13* RNA localization disrupts cell migration and phenocopies acute RAB13 protein loss. **A.** Transwell migration of MDA-MB-231 cells treated with control PMOs or RAB13 PMOs (191+230). Cells reaching the bottom surface after 4hrs were counted. n=25 fields of view in each of 6 independent experiments. **B.** Cells expressing Cherry-NLS, and treated with the indicated PMOs, were tracked every 5min for 10hrs to derive average migration speed. n=65 cells. **C.** PMO-treated cells were induced to invade through a Matrigel plug. Cell staining intensity was used to quantify relative invasion from n=4 independent experiments. **D.** Lifeact-GFP-expressing cells were treated with the indicated PMOs and imaged every minute over 1 hr. Sequential image frames highlight edge retraction (red arrowheads) or protrusion (yellow arrowheads). Corresponding edge velocity is shown, with negative values indicating retraction and positive values indicating extension. Average protrusion and retraction speeds were calculated from n=11-13 cells. **E.** Cells treated with the indicated PMOs or siRNAs were analyzed by western blot to detect RAB13 and GAPDH protein levels. Migration speed was assessed as in (B) from n=55-78 cells. **F.** Protrusion and retraction speed of cells treated with the indicated PMOs or siRNAs was assessed as in (D). Graphs show average normalized values from n=10-13 cells imaged for 1 hr. p-values: *<0.05; **<0.01, ***<0.001, ****<0.0001, by Student’s t-test (A, B, C, D) or analysis of variance with Dunnett’s multiple comparisons test (E, F). n.s.: not significant.

To assess in more detail the molecular mechanisms and polarized behaviors that underlie the observed migration defects, we examined the dynamics of protrusive or retractive cellular regions upon *RAB13* RNA mislocalization. Specifically, Lifeact-GFP expressing cells were imaged over time and the rate of protrusion extension or retraction was quantified (Figure 5D and movie S2). Interestingly, mislocalization of *RAB13* RNA to perinuclear regions significantly reduced the rate of both protrusion and retraction (Figure 5D). Therefore, peripheral localization of *RAB13* RNA appears to impact overall the rate of cytoskeletal dynamics.

### Mislocalization of RAB13 RNA from the periphery phenocopies acute RAB13 protein loss

The cell migration defects observed upon mislocalization of *RAB13* RNA are quite remarkable given that the same amount of RAB13 protein is expressed in the cells (Figure 4C). We thus sought to determine the extent to which the overall RAB13 protein function is compromised when its RNA is prevented from reaching the periphery. For comparison, to set a baseline level, we knocked-down RAB13 protein expression using siRNAs. Indeed, siRNA transfection reduced RAB13 protein to almost undetectable levels (Figure 5E). Consistent with prior reports (Ioannou et al., 2015), RAB13 knockdown resulted in reduced migration speed and compromised the rate of both protrusive and retracting motions (Figure 5E, F). Strikingly, the extent of these defects was very similar to that observed upon *RAB13* RNA mislocalization (Figure 5E, F). Therefore, while translation of *RAB13* RNA in a perinuclear location gives rise to the same amount of RAB13 protein product, the resulting protein is virtually non-functional towards cell migration, phenocopying the effect observed by an almost complete RAB13 protein loss.

### Peripheral RAB13 RNA translation is required for RAB13 protein activation but not steady state distribution or membrane association

To understand this striking effect, we sought to determine how perinuclear versus peripheral translation affects the encoded RAB13 protein. To further confirm that any observed effects are due to changes in *RAB13* RNA distribution, and not due to other potential non-specific consequences of PMO delivery, we additionally examined exogenously expressed GFP-RAB13 protein expressed from a construct that carries the full length, wild-type RAB13 3’UTR or from a construct that carries the same UTR with a deletion of a 52nt region that corresponds to the GA-rich region targeted by the mislocalizing RAB13 PMOs (ΔPMO UTR) (Figure S4A). As predicted, FISH analysis showed that the exogenous wild-type *GFP-RAB13* RNA achieved a peripheral localization, while peripheral localization of the ΔPMO *GFP-RAB13* RNA was significantly abrogated and was similar to the localization of *GFP* RNA (Figure S4A). Exogenous *GFP-RAB13* RNAs were expressed at similar levels (Figure S4B) and did not affect the localization of another endogenous RNA (*NET1*; Figure S4A). Furthermore, the amount of GFP-RAB13 protein produced from the two constructs was indistinguishable, as assessed by Western blot analysis or flow cytometry of GFP expression (Figure S4C, D).

Interestingly, the steady-state localization of RAB13 protein was not impacted upon changing the location of its translation. Endogenous RAB13 protein assumed the same membranous and perinuclear distribution upon PMO delivery (Figure 6A). Similarly, exogenous GFP-RAB13 showed identical distribution regardless of where its RNA was localized (Figure 6B). Because Rab GTPases associate with membranes to control vesicle trafficking, we further examined whether association of RAB13 with membranes was affected by the location of translation. For this, we permeabilized cells with digitonin and isolated soluble and particulate fractions. Again, we found that both endogenous and exogenous RAB13 protein associated similarly with the particulate fraction regardless of where the encoding RNA was enriched (Figure 6C and S5A). Furthermore, consistent with the observed similar association with the membrane fraction, RAB13 protein translated peripherally or perinuclearly associates to the same extent with the Rab escort protein REP-1 (Figure S5B), an interaction required for prenylation of Rab GTPases (Leung et al., 2006a) as well as with RabGDI (Figure S5B), a protein that extracts Rab GTPases from membranes and maintains them soluble in the cytoplasm (Muller and Goody, 2018). Therefore, despite the pronounced functional defect exhibited upon *RAB13* RNA mislocalization, at least some of the properties and interactions of RAB13 protein are not affected.

**Figure 6:**
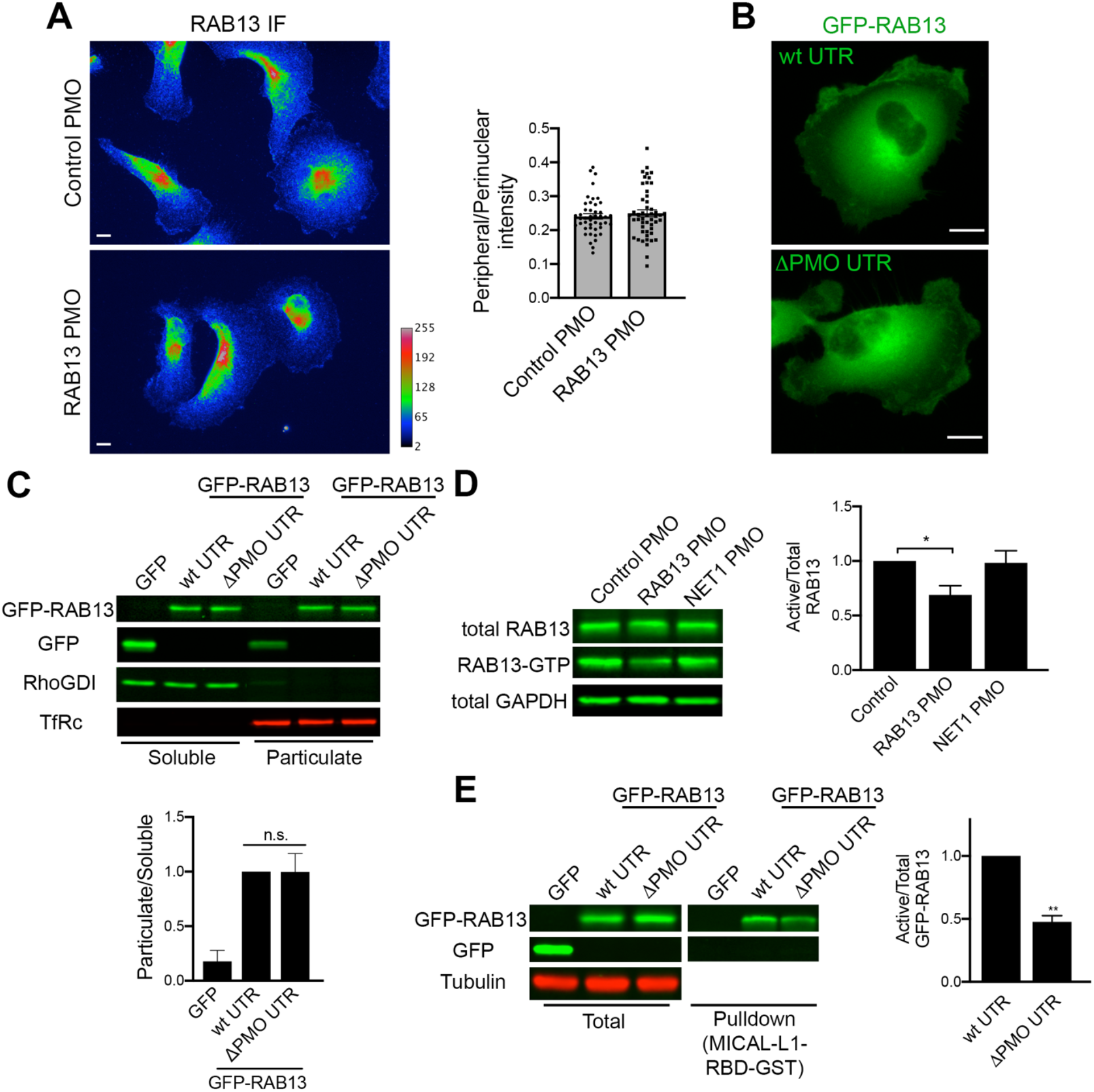
Peripheral *RAB13* RNA translation is required for RAB13 protein activation but not steady state distribution or membrane association. **A.** RAB13 immunofluorescence of MDA-MB-231 cells treated with control or RAB13 (191+230) PMOs and ratios of peripheral/perinuclear intensity. n=45-50 cells. Similar results were observed in two additional independent experiments. **B.** Fluorescence images of cells expressing GFP-RAB13 with the indicated UTRs. Note that in both cases the protein assumes indistinguishable distribution. Scale bars: 10 μm. **C.** Soluble/particulate fractionation of the indicated cell lines followed by Western blot to detect the indicated proteins. RhoGDI and TfRc serve as soluble and particulate markers, respectively. Graph shows quantitation from n=3 independent experiments. **D, E.** Active RAB13 (RAB13-GTP) was pulled-down using MICAL-L1 RBD-GST from the indicated PMO-treated cells (D) or GFP-RAB13 expressing lines (E). The amount of endogenous or exogenous RAB13 was measured by quantitative Western and relative levels of active RAB13 are plotted. n=8 (D), n=4 (E). p-values: *<0.05, **<0.01 by Kruskal-Wallis test.

We then investigated whether perinuclear translation affects RAB13 activity. Active Rab proteins are loaded with GTP through the action of guanine nucleotide exchange factors (GEFs). GTP-loading promotes Rab binding to effector molecules. We assessed RAB13 activity through measuring its ability to interact with the RAB13-binding domain (RBD) of its effector protein MICAL-L1 (Ioannou et al., 2015). Interestingly, endogenous RAB13 produced from a perinuclear RNA (RAB13 PMO cells) exhibited significantly reduced activity compared to peripherally translated RAB13 protein (control PMO cells or cells treated with PMOs against the unrelated *NET1* RNA) (Figure 6D). In agreement, pulldown assays with the MICAL-L1 RBD also revealed that the exogenous GFP-RAB13 protein expressed from a perinuclearly localized RNA had significantly less activity compared to peripherally translated protein (GFP-RAB13 with wtUTR) (Figure 6E). Therefore, RAB13 activity and function in cell migration is determined by the subcellular location of *RAB13* RNA translation.

### Peripheral RAB13 RNA promotes the local association of RAB13 with the exchange factor RABIF

To understand the underlying mechanism, we used mass spectrometry analysis to identify RAB13-interacting proteins. Specifically, we identified proteins bound to wild-type GFP-RAB13 or GFP-RAB13(T22N), a nucleotide-free mutant which binds tightly to putative GEFs (Table S1). As expected, wild-type GFP-RAB13 associated with RabGDI isoforms. The GFP-RAB13(T22N) mutant did not associate with RabGDI but interacted with RABIF/MSS4, a protein known as a GEF for Rab GTPases (Burton et al., 1993, Burton et al., 1994, Miyazaki et al., 1994) (Table S1). Western blot analysis verified the association of RABIF with RAB13 and revealed that it interacts preferentially with RAB13(T22N), as would be expected from a GEF (Figure 7A, B). To confirm that RABIF acts as a RAB13 GEF in our system, we used CRISPR/Cas9 to knockout its expression (Figure 7C). Indeed, two different guide RNAs targeting RABIF led to a significant reduction of RAB13 activation levels (Figure 7C). Furthermore, importantly, expression of RAB13(T22N) from a perinuclear RNA (carrying the ΔPMO UTR) led to a significantly reduced association with RABIF (Figure 7 A, B), strongly suggesting that RABIF is part of the mechanism linking *RAB13* RNA localization to RAB13 protein activation.

**Figure 7:**
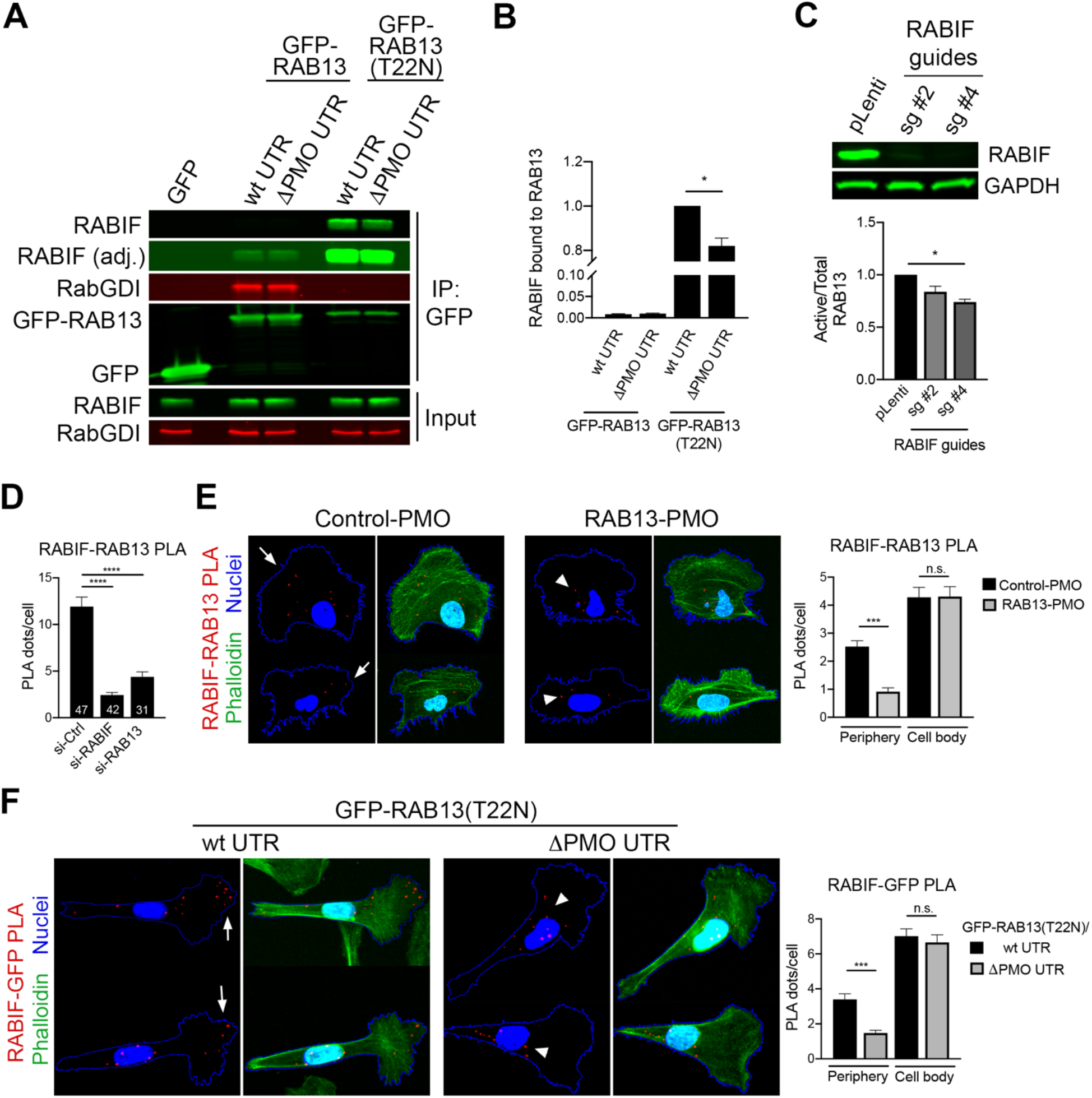
Peripheral *RAB13* RNA promotes the local association of RAB13 with the exchange factor RABIF. **A.** Immunoprecipitation and Western blot analysis to detect proteins bound to GFP-RAB13 expressed from constructs carrying wt or ΔPMO RAB13 3’UTR. RABIF panel is also shown with adjusted contrast to reveal lower binding to wtGFP-RAB13. **B.** Quantification of RABIF binding to RAB13 from experiments as in (A). n=6. **C.** Active RAB13 pulldown assay from cells with CRISPR knockdown of RABIF using the indicated sgRNAs. n=3. **D.** Quantification of RABIF-RAB13 PLA signal from cells transfected with the indicated siRNAs. Number of observed cells is indicated within each bar. **E.** Representative RABIF-RAB13 PLA images and quantitations from cells transfected with the indicated PMOs. N>70 cells. **F.** Representative RABIF-GFP PLA images and quantitations from cells expressing GFP-RAB13(T22N) carrying wt or ΔPMO RAB13 3’UTR. N>70 cells. Arrows indicate PLA signal at the cell periphery. Arrowheads indicate PLA signal within the cell body. p-values: *<0.05, **<0.01, ***<0.001, ****<0.0001 by Wilcoxon matched-pair signed rank test (B) or analysis of variance with Dunn’s, Dunnett’s or Tukey’s multiple comparisons test (C-F).

The above data show that peripheral localization of *RAB13* RNA affects the degree of RAB13-RABIF interaction. However, both proteins exhibit a steady-state perinuclear enrichment (Figure 1B and S6A). To understand this effect, we examined whether this interaction occurs in a spatially-specific manner. For this, we used proximity ligation assay (PLA) to detect protein-protein interactions *in situ*. PLA between endogenous RAB13 and RABIF revealed a specific interaction between the two, since the signal was significantly decreased upon knockdown of each binding partner (Figure 7D and S7A). RAB13-RABIF interaction was detected in both peripheral and perinuclear regions. However, interestingly, mislocalization of *RAB13* RNA upon PMO delivery, reduced specifically the peripheral RAB13-RABIF interaction (Figure 7E and S7B). Similarly, interaction between GFP-RAB13(T22N) and RABIF occurred in both peripheral and perinuclear regions and expression of GFP-RAB13 from a mislocalized construct (carrying the ΔPMO UTR) specifically reduced the observed peripheral interaction (Figure 7F). Thus, a peripheral pool of RAB13-RABIF complex requires peripheral localization of *RAB13* RNA.

### RABIF associates with RAB13 co-translationally

The above data suggest that RABIF might interact with RAB13 co-translationally and thereby where *RAB13* RNA is located and translated determines the site of complex formation. To determine whether a co-translational interaction occurs, we immunoprecipitated RABIF and tested for the presence of *GFP-RAB13* RNA (Figure 8A). Indeed, both *GFP-RAB13* and *GFP-RAB13(T22N)* RNAs were readily detected above background in RABIF immunoprecipitates (Figure 8B). To further assess whether this interaction depends on active translation, immunoprecipitations were carried out from cells pre-treated with puromycin, an antibiotic which dissociates translating ribosomes. Importantly, puromycin treatment abolished the binding of RABIF to *RAB13* RNA (Figure 8B). These data indicate that RABIF does not directly associate with *RAB13* RNA. We rather interpret these data to indicate that RABIF co-translationally binds to the nascent RAB13 protein.

**Figure 8:**
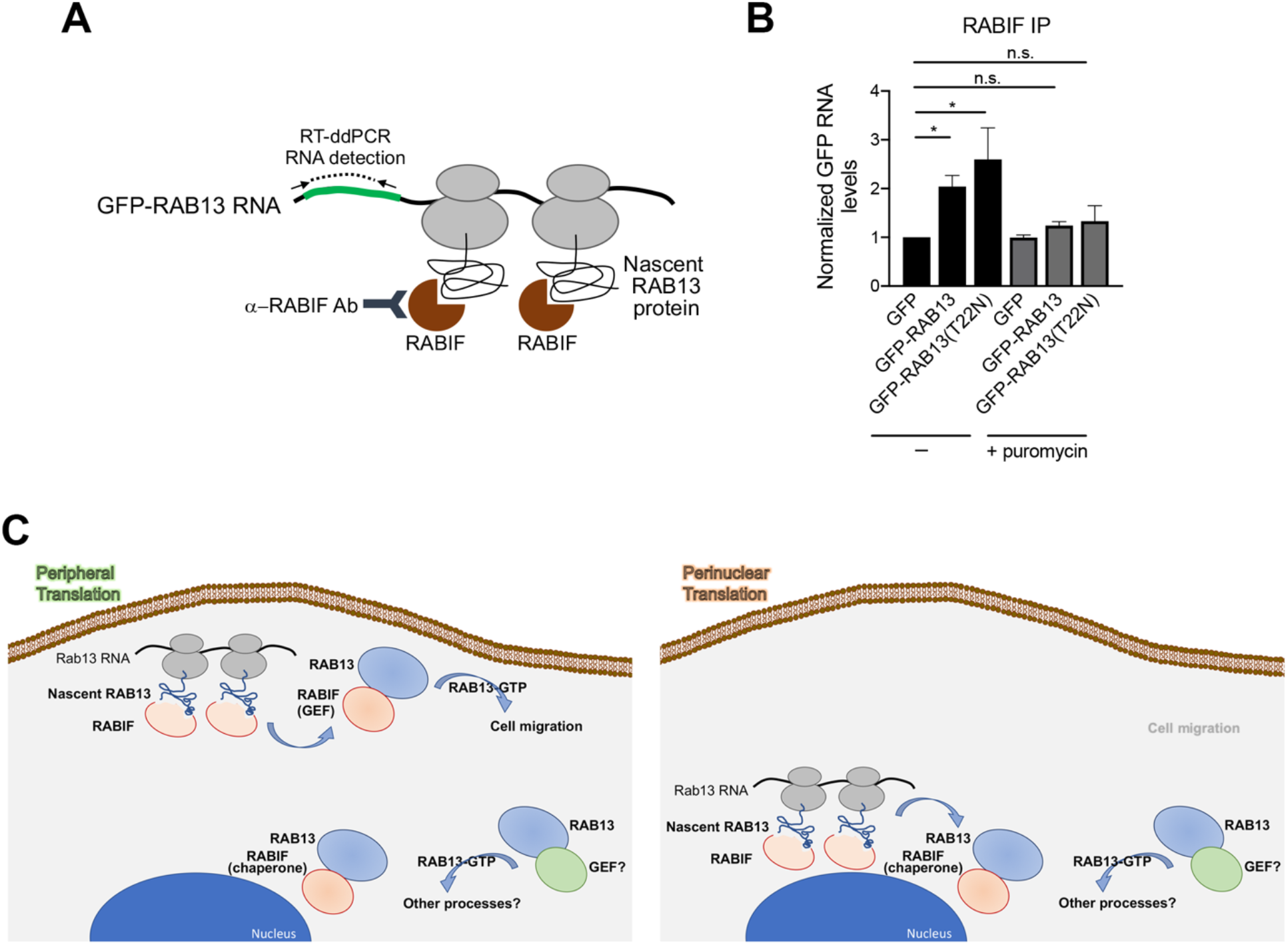
RABIF associates with RAB13 co-translationally. **A.** Schematic depicting experimental strategy for assessing co-translational association of RABIF with nascent RAB13. **B.** Quantification of GFP RNA levels associating with RABIF in immunoprecipitation assays from the indicated cell lines. Note that even though RABIF binds several-fold more to RAB13(T22N) (see Figure 7A, B) it binds similarly to the wild-type and T22N *RAB13* RNA, indicating that RABIF binds similarly to nascent RAB13. After translation, it is likely displaced upon GTP-loading of wild type RAB13, while it remains more stably bound to the nucleotide-free (T22N) form. N=6. p-values: *<0.05 by analysis of variance with Dunn’s multiple comparisons test. **C.** Proposed model for regulation of RAB13 activity through local RNA translation. See text for details.

## DISCUSSION

We show here that, for the protrusion-localized *RAB13* RNA, peripheral translation does not lead to a corresponding local protein accumulation. Rather the location of translation critically affects some, but not all properties of the encoded RAB13 protein. Specifically, the location of translation does not affect the steady-state RAB13 protein distribution, its membrane association or interaction with Rab regulators such as REP-1 and RabGDI. Strikingly however, translation at the periphery is required for full RAB13 activation through a co-translational association with the RABIF GEF (Figure 8C). Preventing peripheral translation compromises the ability of RAB13 to support cell migration to an extent similar to that observed upon almost complete loss of the protein. This work identifies the subcellular location of *RAB13* RNA translation as a critical factor determining the exact functionality and properties of the encoded protein.

It is increasingly appreciated that the process of translation can affect the encoded proteins through various mechanisms. For instance, the translation machinery provides a platform for recruitment of maturation factors that guide nascent polypeptide chains into functional protein structures (Gloge et al., 2014). Furthermore, subunits of hetero-oligomeric complexes come into contact and begin assembling co-translationally (Shiber et al., 2018). Such co-translational interactions can determine the efficiency of functional complex formation (Shieh et al., 2015). Additionally, the untranslated regions (UTRs) of the RNA template can impact the fate of the newly synthesized proteins by acting as scaffolds to recruit protein complexes that affect the targeting or modification of the nascent or newly-synthesized polypeptides (Basu et al., 2011, Berkovits and Mayr, 2015). In this way, use of alternative UTRs can direct formation of different protein complexes that support distinct downstream protein functions (Berkovits and Mayr, 2015, Lee and Mayr, 2019). Our results suggest that another factor that can have a determining impact on downstream protein activity is the subcellular microenvironment into which a protein is being synthesized.

Peripheral translation affects RAB13 function without resulting in a corresponding accumulation of RAB13 at the periphery. This finding contrasts with observations, in yeast or mammalian neurons, of other localized RNAs whose translation leads to local increase in protein concentration (Holt et al., 2019, Paquin and Chartrand, 2008, Rangaraju et al., 2017). A common feature of these latter cases is that premature or ectopic translation would have deleterious consequences, such as disruption of cell type identity in yeast, or incorrect axonal pathfinding or synaptic marking in neuronal cells. We believe that another relevant point is that such examples of localized RNA transcripts, for the most part, pertain to RNAs which are localized in rather stably polarized structures, whose identity remains fixed even though specific signals might elicit transient responses. By contrast, protrusions of migrating cells are very dynamic switching between extension and retraction phases as they explore the surrounding environment (Ryan et al., 2012). Directional migration requires that cells are able to quickly repolarize and redefine their leading and retracting areas to follow the direction of an asymmetric extracellular cue (Petrie et al., 2009). Given that such dynamic changes can occur within minutes, which is the same range as the time required for translation of an average size protein, it seems unlikely that the use of local translation as a means of building dynamic local concentration gradients would be an efficient process. In light of the evidence presented here, we rather propose that in dynamically changing protrusions local translation serves to expose newly synthesized proteins to the local environment and thus impart on them particular properties that might persist even after they have diffused or trafficked away from their site of synthesis. Given that global studies have shown little correlation between RNAs localized at cell protrusions and steady-state concentration of the corresponding proteins (Mardakheh et al., 2015), it would appear that this mode of regulation might be relevant for the majority of protrusion-localized RNAs.

For RAB13, peripheral translation is required for its association with RABIF in this local environment. Our data indicate that co-translational RABIF-RAB13 association at the periphery is required for full RAB13 activation and for RAB13 function in cell migration. We propose that this peripherally-activated RAB13 pool is specifically directed towards migration-relevant effectors, since mislocalization of *RAB13* RNA phenocopies complete loss of protein expression with regards to cell migration (Figure 8C). Interestingly, this occurs even though a substantial amount of GTP-loaded, membrane-bound RAB13 can still be detected. Therefore, RAB13 protein, that is not produced peripherally, can be activated and likely contributes to different cellular functions, perhaps by interacting with different effectors. Indeed, RAB13 has been shown to participate in diverse trafficking events including the recycling of integrins (Ioannou et al., 2015, Nishikimi et al., 2014, Sahgal et al., 2019, Wu et al., 2011) or of factors involved in cell-cell contacts (Kohler et al., 2004, Marzesco et al., 2002, Morimoto et al., 2005, Terai et al., 2006). Given that significant RAB13 activity remains in the absence of RABIF, we favor the idea that RAB13 activation at perinuclear locations relies on other GEFs. These could include proteins of the DENND family, such as DENND2B and DENND1C (Ioannou et al., 2015, Nishikimi et al., 2014) even though we haven’t been able to detect expression of DENND2B in MDA-MB-231 cells. RABIF additionally exhibits chaperone activity (Gulbranson et al., 2017) and consistent with that we observe a reduction of RAB13 levels upon RABIF knockout (Figure S6B), but not upon *RAB13* RNA mislocalization. We propose that RABIF affects RAB13 differently based on the particular subcellular environment, acting as a GEF at the periphery and as a chaperone in the cell body (Figure 8C). Overall, our data highlight a role for local translation in directing Rab GTPase activation and suggest that the micro-environment into which a protein is synthesized can affect its regulation and downstream function.

## ACKNOWLEDGEMENTS

We would like to thank the CCR Genomics Core at National Cancer Institute/NIH for help with ddPCR and the NCI LGI Flow Cytometry Core for help with cell sorting and flow cytometry. This work was supported by the Intramural Research Program of the Center for Cancer Research, NCI, National Institutes of Health (S.M.; 1ZIA BC011501).

## AUTHOR CONTRIBUTIONS

Conceptualization, S.M. and K.M.; Methodology, S.M., K.M., M.L.H.; Formal analysis, K.M., M.S., A.N.G., T.W. L.M.J. and S.M.; Investigation, K.M., M.S., A.N.G., T.W. and S.M.; Writing-original draft; K.M. and S.M.; Writing-review and editing, K.M., A.N.G., T.W., M.L.H. and S.M.

## MATERIALS AND METHODS

### Plasmid constructs and lentivirus production

To express β-globin RNA followed by different UTRs, the genomic region containing the coding sequence of human β-globin was ligated to fragments corresponding to the following UTRs: β-globin UTR (3′UTR of human β-globin, Acc#: NM_000518.4), Rab13 UTR (wt) (3′UTR of mouse Rab13, Acc#: NM_026677.4), Rab13 UTR (Δ1) (3′UTR of mouse Rab13 missing nucleotides 204-211, corresponding to region 1), and Rab13 UTR (Δ1+2) (3′UTR of mouse Rab13 missing nucleotides 204-211 and 193-200, corresponding to regions 1 and 2, respectively). The sequences were cloned into the pENTR1A vector and then transferred into the pINDUCER20 lentivector (Addgene #44012) using the Gateway LR clonase II Enzyme mix (Thermo Fisher Scientific, cat# 11791-020).

mEGFP-Lifeact-7 (gift of Michael Davidson; Addgene plasmid # 54610) was transferred into pCDH-CMV-MCS-EF1-Puro (System Biosciences, cat #CD510B-1) using NheI/NotI sites for virus production.

To express N-terminally GFP-tagged RAB13 using different UTRs, the coding sequence of EGFP was ligated in frame to the coding sequence of human RAB13. The fusion was then ligated to fragments corresponding to the 3’UTR of human RAB13 (Acc#: NM_002870.5) (wt UTR) or the 3’UTR of RAB13 missing 53 nucleotides (202-254) from the GA-rich region that is targeted by the RAB13 PMOs (ΔPMO UTR). The sequences were cloned into PCDH-PGK lentivector.

Lentivirus was produced in HEK293T cells cultured in DMEM containing 10% FBS and Penicillin/Streptomycin. HEK293T cells were transfected with pINDUCER20 lentivectors, together with packaging plasmids pMD2.G and psPAX2 using PolyJet In Vitro DNA transfection Reagent (SignaGen) for 48 h. Harvested virus was precipitated with Polyethylene Glycol at 4 °C overnight.

### Immunofluorescence and Western blot

For IF, cells were plated onto collagen IV-coated glass coverslips (10μg/ml) and then fixed with 4% paraformaldehyde (PFA) in PBS (phosphate-buffered saline) for 15min, permeabilized with 0.2% Triton X-100 in PBS for 5 min, blocked with 5% goat serum in PBS for 1h, and incubated with Rab13 rabbit polyclonal antibody (Novus Biologicals, cat# NBP1-85799, 1/200 dilution) for 1h. Secondary antibodies were conjugated with Alexa 647 (Thermo Fisher Scientific).

For Western blot detection the following antibodies were used: anti-Rab13 rabbit polyclonal (Novus Biologicals, cat# NBP1-85799, 1/1,000 dilution), anti-GFP rabbit polyclonal (Invitrogen, cat# A11122, 1/2,000 dilution), anti-RhoGDI, anti-Transferrin receptor (clone H68.4, ThermoFisher Scientific, cat# 13-6800), anti-α-tubulin mouse monoclonal (Sigma-Aldrich, cat# T6199, 1/10,000 dilution), and anti-GAPDH rabbit monoclonal 14C10 (Cell Signaling Technology, cat# 2118, 1/2,000 dilution), anti-phospho-FAK(Y397) (Cell Signaling Technology, cat# 3283), anti-REP-1 (Abcam, cat# 134964), anti-RabGDI (Santa Cruz Biotechnology, cat# sc-374649), anti-RABIF mouse monoclonal antibody D-12 (Santa Cruz biotechnology, cat# sc-390759, 1/1,000 dilution).

### Cell culture, siRNA transfection and generation of stable cell lines

NIH/3T3 mouse fibroblast cells (ATCC) were grown in DMEM supplemented with 10% calf serum, sodium pyruvate and penicillin/streptomycin (Invitrogen) at 37°C, 5% CO_2_. MDA-MB-231 human breast cancer cells (ATCC) were grown in Leibovitz’s L15 media supplemented with 10% fetal bovine serum and penicillin/streptomycin at 37°C in atmospheric air. Cells have been tested for mycoplasma and are free of contamination.

For knockdown experiments, 40 pmoles of siRNAs were transfected into cells with Lipofectamine RNAiMAX (Thermo Fisher Scientific, cat# 13778-150) according to the manufacturer’s instructions. Cells were assayed after three days. siRNAs used were: AllStars Negative control siRNA (cat# 1027281), si-Rab13 #7 (cat# SI02662149; target sequence: 5’- CAGGGCAAACATAAATGTAAA-3’), si-Rab13 #8 (cat# SI02662702; target sequence: 5’- ATGGTCTTTCTTGGTATTAAA-3’), si-RABIF (Cat# SI00301595) from Qiagen.

To generate stable cell lines expressing β-globin reporters with different UTRs, NIH/3T3 cells were infected with the corresponding lentiviruses and selected with 0.6mg/ml Geneticin (Thermo Fisher Scientific). Exogenous expression of the reporters was induced by addition of 1 μg/ml Doxycycline (Fisher Scientific) approximately 2-3 hrs before processing cells for FISH. This concentration of Doxycycline and duration of incubation was chosen to achieve relatively low levels of expression and prevent competition effects.

To generate stable cell lines expressing GFP-RAB13 with different UTRs, MDA-MB-231 cells were infected with the corresponding lentiviruses and selected with 6μg/ml Blasticidin (Thermo Fisher Scientific). GFP-expressing cells with low level of GFP expression were sorted by FACS.

To generate stable RABIF knockout cell lines using CRISPR-Cas9 technology, sgRNAs targeting RABIF (sg #2: 5’-GAGCGAGTTAGTGTCAGCCGAGG-3’ and sg #4 5’-AGCGAGTTAGTGTCAGCCGAGGG-3’) were cloned in lentiCRISPRv2 (Addgene #52961). MDA-MB-231 cells were infected with the corresponding lentiviruses and selected with 1μg/ml puromycin (Thermo Fisher Scientific).

### Morpholino ASOs and delivery

PMOs were synthesized by Gene Tools, LLC and delivered into cells using EndoPorter(PEG) (GeneTools, LLC). Sequences used are listed in Table S2.

### Protrusion/cell body isolation and RNA analysis

Protrusions and cell bodies were isolated from serum-starved cells plated for 2 hrs on Transwell inserts equipped with 3.0-μm porous polycarbonate membrane (Corning) as previously described (Wang et al., 2017) with some modifications. Briefly, 3.5 million serum-starved cells were plated per 25mm filter, pre-coated with 10 μg/ml collagen IV, and 1 or 3 filters were used for cell body or protrusion isolation, respectively. Cells were allowed to attach for 2hrs and LPA (150ng/ml) was added to the bottom chamber for 40min. The cells were fixed with 0.3% paraformaldehyde for 10 min. For isolating protrusions, cell bodies on the upper surface were manually removed by wiping with cotton swab and laboratory paper. The protrusions on the underside were then solubilized by immersing the filter in crosslink reversal buffer (100mM Tris pH 6.8, 5mM EDTA, 10mM DTT and 1% SDS) and gentle scraping. Cell bodies were similarly isolated after manually removing protrusions from the underside of the membrane. The extracts were incubated at 70 °C for 45 min and used for protein analysis or RNA isolation using Trizol LS (Thermo Fisher Scientific).

For nanoString analysis, RNA samples were analyzed using a custom-made codeset and the nCounter analysis system according to the manufacturer’s instructions (NanoString Technologies). For ddPCR, RNA samples were analyzed using the ddPCR EvaGreen Supermix (Bio-Rad, cat. no. 186-4034). Droplets were generated using the Automated Droplet Generator (Bio-Rad, cat no. 186-4101), PCR amplification was performed on a C1000 Touch™ Thermal Cycler (Bio-Rad, cat no. 185-1197) and droplet reading was done with the QX 200 Droplet reader (Bio-Rad, cat no. 186-4003) and QuantaSoft software (Bio-rad)

### Fluorescence in situ hybridization (FISH)

For FISH, cells were plated on fibronectin- or collagen IV-coated glass coverslips for 2-3 h and subsequently fixed with 4% PFA for 20min (5μg/ml fibronectin for NIH/3T3 cells and 10μg/ml collagen IV for MDA-MB-231 cells). FISH was performed with ViewRNA ISH Cell Assay kit (Thermo Fisher Scientific) according to the manufacturer’s instructions. The following probe sets were used: human HBB #VA1-13382; mouse Ddr2 #VB6-12897; mouse Rab13 #VB1-14374; mouse Cyb5r3 #VB6-3197970; human Rab13 #VA1-12225; human Net1 #VA6-3169338; GFP #VF6-16198. DAPI was used to stain nuclei and CellMask stain (Thermo Fisher Scientific) was used to identify the cell outlines. Samples were mounted with ProLong Gold antifade reagent (Thermo Fisher Scientific). Image analysis and quantification of RNA distributions was performed using RDI Calculator (Stueland et al., 2019).

### Active RAB13 pull-down assays, cell fractionation and immunoprecipitation

For RAB13 pull-down, cells were washed with ice-cold TBS and lysed in buffer containing 50 mM Tris (pH 7.4), 150 mM NaCl, 10 mM MgCl_2_, 1% Triton X-100, 0.1% SDS, 0.5% sodium deoxycholate, 10% glycerol and protease inhibitor cocktail. Clarified lysates were then incubated with 20 μg of recombinant glutathione *S*-transferase (GST)-RBD-MICAL-L1 and Pierce™ Glutathione magnetic agarose beads (Life Technologies-Invitrogen, cat# 78601) for 2 hrs at 4°C. Beads were washed with lysis buffer without sodium deoxycholate and SDS, eluted with sample buffer, and analyzed by immunoblotting.

To obtain soluble and particulate fractions, adherent cells were incubated with a buffer containing 50mM Tris pH 7.4, 50mM NaCl, 0.01% Digitonin, 10mM MgCl_2_, 1mM EDTA, 10% glycerol and a cocktail of protease and phosphatase inhibitors (Thermo Fischer Scientific, cat# 1861281) with rocking at 4°C for 10min. Cell material extracted in this buffer was defined as soluble. The remaining material, corresponding to the particulate fraction, was extracted using a buffer containing 50mM Tris pH 7.4, 1% Triton X-100, 150mM NaCl, 10mM MgCl_2_, 0.1% SDS, 0.5% deoxycholate and 10% glycerol. Equal volumes of these fractions were analyzed by SDS-PAGE and immunoblotting. For quantitation, the soluble and particulate amounts were normalized to the corresponding amounts of soluble and particulate markers.

For immunoprecipitations, cells were lysed with a buffer containing 50mM Tris pH 7.4, 1% NP-40, 150mM NaCl, 10mM MgCl_2_, 10% glycerol and a cocktail of protease and phosphatase inhibitors (Thermo Fischer Scientific, cat# 1861281). Lysates were cleared by centrifugation and mixed with GFP-Trap Magnetic Agarose beads (Chromotek, cat# gtma-10) or protein G Dynabeads (Fisher Scientific) for 1.5 h, at 4°C. Immobilized complexes were eluted with Laemmli’s buffer and analyzed by SDS-PAGE and immunoblotting.

### Migration assays

Transwell migration assays were performed using 24-well plates with 8μm-pore size inserts. Inserts were coated on both sides with collagen IV overnight (10μg/ml in PBS). MDA-MB-231 cells that had been kept in 0.1% FBS-containing media for 16 h were detached using trypsin. After addition of trypsin inhibitor cells were resuspended in media containing 0.1% FBS and 20,000 cells (in 400μl) were plated onto each insert. The lower chamber contained 600μl of the same media. 2 h post-plating, 10% FBS was added in the lower chamber to induce cell motility. 2 h later both sides of the membrane were fixed with 4% PFA, cells were removed from the upper chamber and the membrane was stained with DAPI. Technical replicates were included in each assay. Nuclei were counted within 25 fields of view in each insert (5×5 tiles; tile area=456μm×456μm).

Invasion assays were performed based on (Scott et al., 2011) with some modifications. 24-well plates with 8 μm pore size inserts were used. Matrigel (BD Biosciences, cat# 354324) was used at 50% dilution with PBS. 100μl of matrix solution was added to the inserts and let to solidify. 40,000 cells (in 100μl regular media) were plated on the upward facing underside and allowed to attach for 4 h. Subsequently, transwells were inverted, the underside was washed with serum-free media and transwells were transferred in wells containing 500μl of serum-free media. 100μl of 10% FBS-containing media were added on top of the solidified Matrigel and cells were allowed to invade for 48 h. Transwells were fixed in 4% PFA/0.2% Triton X-100 in PBS for 1hr at room temperature and nuclei were stained with DAPI overnight.

### Proximity ligation assay

Cells plated on Collagen IV (Sigma, Cat# C5533)-coated coverslips were fixed with 4% formaldehyde in PBS for 15 min and then permeabilized with 0.5% Triton X-100 in PBS for 5 min at room temperature. Cells were subsequently blocked in Duolink blocking solution (Sigma Aldrich, cat# DUO92101) for 1h at 37C and incubated with a pair of primary antibodies diluted in Duolink antibody diluent (in a humidified chamber for 1.5h at room temperature). Antibodies used were anti-Rab13 (1:200; Novus Biologicals, cat# NBP1-85799), anti-GFP (1:8,000; Invitrogen, cat# A11122) and anti-RABIF (1:1,500; Santa Cruz biotechnology, cat# sc-390759). After washing, PLA probes were applied in 1:10 dilution using the diluent buffer provided in the Duolink In Situ Red kit (Sigma Aldrich, cat# DUO92101). Incubations and subsequent ligation and amplification steps were performed according to the manufacturer’s instructions. After the final washes, cells were post-fixed for 10 min at room temperature with 4% formaldehyde in PBS, stained with Alexa Fluor-488 phalloidin (Thermo Fisher, cat# A12379) in blocking buffer for 30 min and mounted using Duolink PLA Mounting medium with DAPI.

### Co-translational association of RABIF with Rab13 RNA

Cells treated with vehicle or 50μg/ml puromycin for 15min (to disrupt sites of translation) were lysed as described above and processed for RABIF immunoprecipitation using protein G Dynabeads (Fisher Scientific). After multiple washes, immune complexes were eluted in the same buffer supplemented with 1% SDS by rocking at RT for 15min. Total RNA present in the eluate or the extract (input) was isolated using Trizol LS (Thermo Fisher Scientific) and reverse transcribed using the iScript cDNA synthesis kit (Biorad). Fixed amount of spike RNA (β-globin) was added in each sample to correct for potential losses during these steps. The abundance of GFP, GAPDH and b-globin RNA was then quantified using digital droplet PCR (Biorad). Values in each eluate were normalized to the corresponding values in the input and the enrichment of GFP-RAB13 RNA over GAPDH was assessed.

### Imaging and image analysis

Images of all fixed samples were obtained using a Leica SP8 confocal microscope, equipped with a HC PL APO 63x oil CS2 objective. Z-stacks through the cell volume were obtained and maximum intensity projections were used for subsequent analysis. Image analysis was performed using ImageJ or RDI Calculator (Stueland et al., 2019).

For live imaging of Lifeact-GFP-expressing cells, cells were plated on collagen IV-coated coverglass and were imaged on a Leica SP8 confocal microscope equipped with HC PL APO 63x oil CS2 objective, at constant 37C temperature and atmospheric air. The 488nm laser line was used for illumination and z-stacks through the cell volume were acquired every 1.5 min over a period of 1hr. Maximal intensity projections were produced and edge velocity was determined using the ADAPT ImageJ plugin (Barry et al., 2015) by thresholding the image and using the plugin’s default settings. Protrusion and retraction velocities were averaged for each cell.

For live imaging of cherry-NLS-expressing cells, cells were plated on collagen IV-coated coverglass and were imaged on an Olympus IX81 microscope equipped with a 10x phase UPlan Fluorite Dry objective, a pE-300ultra Illumination system, an ORCA-Flash4.0 v3 sCMOS camera and an Okolab cage incubator enclosure. Time lapse images were taken every 5min over a period of 10 hours. Nuclei were tracked using the built-in Manual Tracking ImageJ plugin. The resulting XY coordinates for each cell track were imported into a DiPer macro (Gorelik and Gautreau, 2014) to calculate average cell speed.

## SUPPLEMENTARY FIGURES AND FIGURE LEGENDS

**Figure S1:**
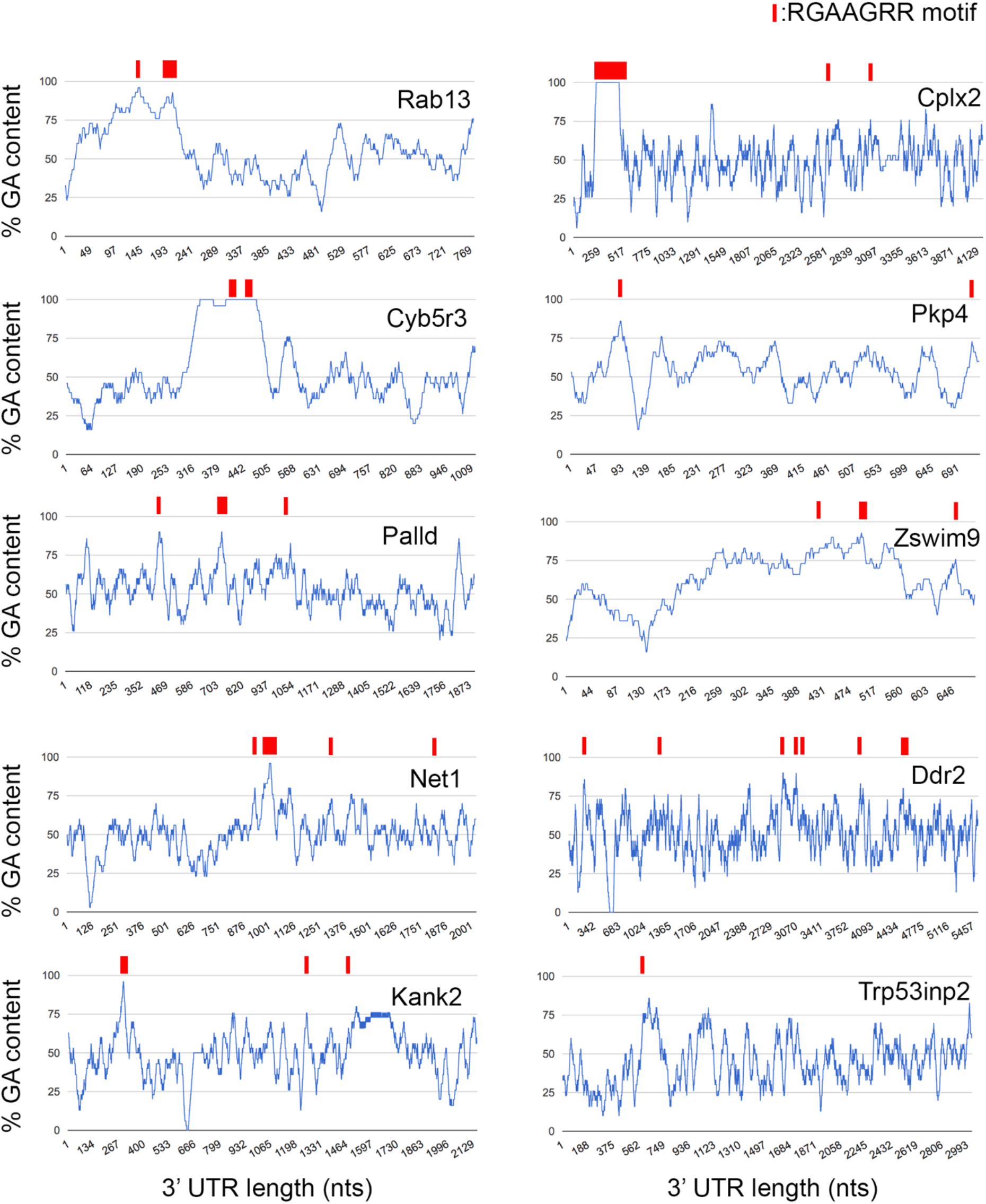
GA-rich motif distribution in 3’UTRs of APC-dependent RNAs. Graphs show the %GA content along the 3’UTR of the indicated APC-dependent RNAs using a 30nt window size. Occurrences of the consensus GA-rich motif are indicated by a red rectangle. Wider rectangles indicate the presence of multiple motifs. Exact sizes are not to scale due to the variable UTR lengths. The majority of GA-rich motifs are found within more extended GA-rich regions with GA content >75%. Note that the GA-motif is 7 nts while the window for %GA calculation is 30 nts.

**Figure S2:**
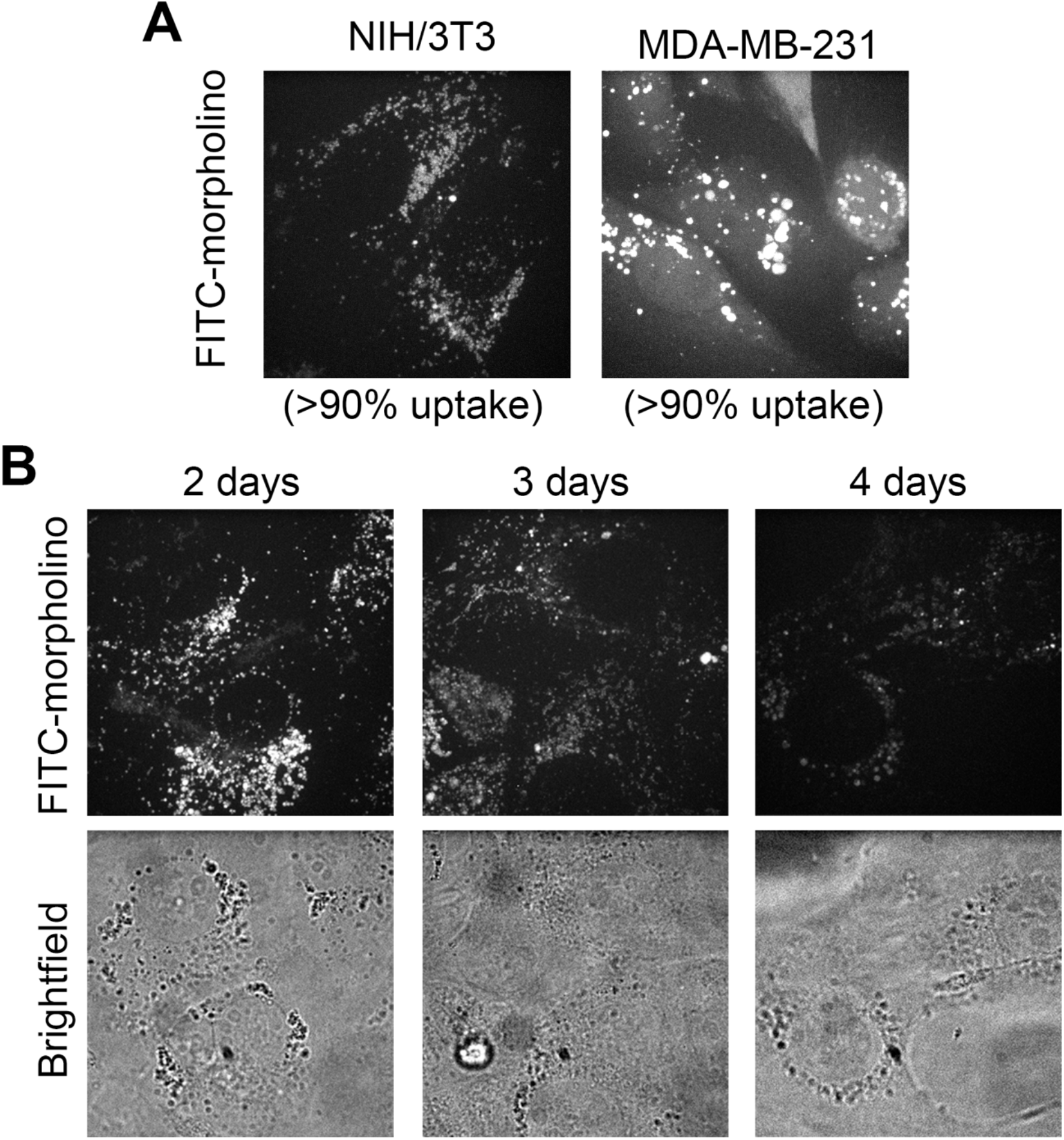
Uptake and persistence of PMOs. **A.** FITC-labeled morpholinos were delivered into the indicated cell lines and fluorescence uptake was assessed after 24 hrs by live cell imaging. >90% of cells had visible fluorescence in intracellular vesicles or diffusely in the cytoplasm. **B.** Mouse fibroblast cells were treated with FITC-labeled morpholinos. Persistence of morpholinos in cells was assessed daily for 4 days by live-cell imaging. Note that after 4 days, signal is still detectable in >90% of cells.

**Figure S3:**
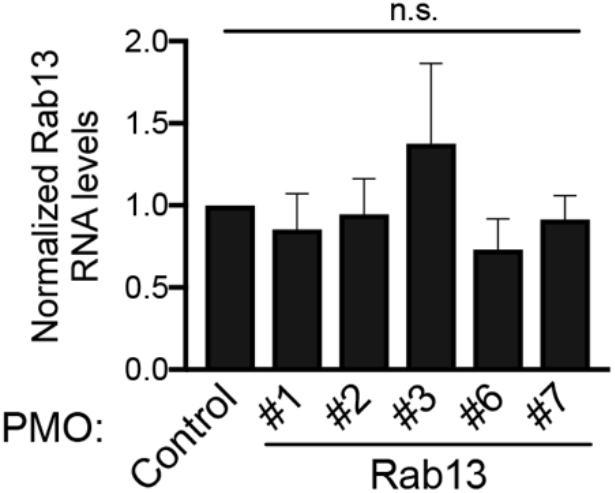
Rab13 PMOs do not affect *Rab13* RNA levels. The indicated PMOs were delivered into mouse fibroblast cells. *Rab13* RNA levels were assessed by RT-ddPCR and normalized to housekeeping RNA levels. n.s.: not significant by one-way ANOVA.

**Figure S4:**
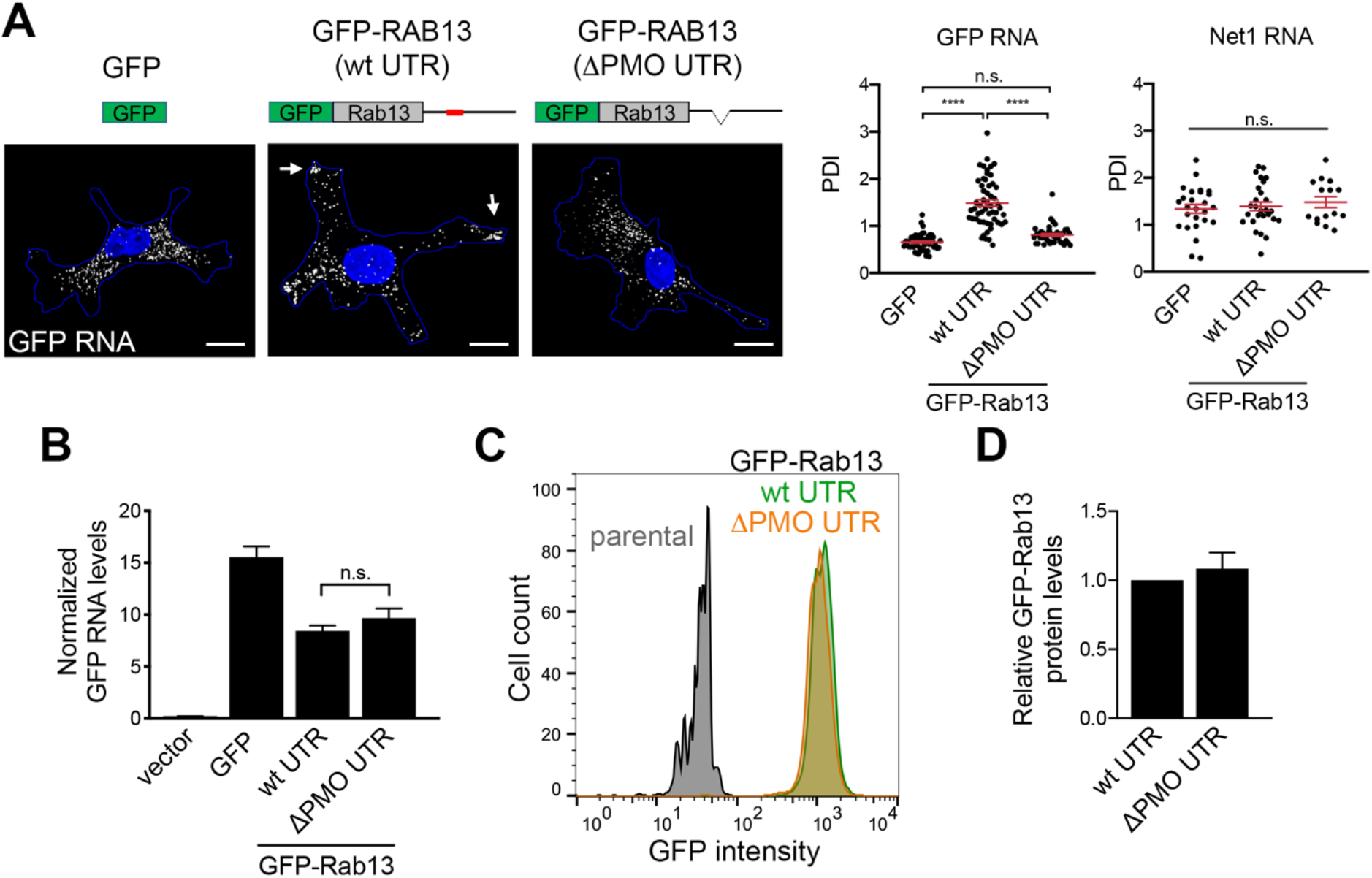
Peripheral localization of exogenous *RAB13* RNA does not affect *RAB13* RNA stability or translation. **A.** Schematics depict GFP or GFP-RAB13 constructs stably expressed in MDA-MB-231 cells. RAB13 coding sequence is followed either by the wild type RAB13 UTR or the RAB13 UTR carrying a 52 nt deletion corresponding to the region targeted by PMOs 191 and 230 (ΔPMO UTR). Exogenous RNA is detected by FISH against the GFP sequence. Arrows point to RNA localized at protrusions. Scale bars: 10 μm. Graphs show PDI measurements of *GFP* or *NET1* RNA from multiple cells. n=42-54 cells in 4 independent experiments (for GFP); n=27 cells in 2 experiments (for Net1). ****: p<0.0001 by analysis of variance with Dunnett’s multiple comparisons test. **B.** Levels of *GFP* RNA from the indicated cells lines were assessed by RT-ddPCR and normalized to housekeeping control RNAs. n=6. **C.** The indicated cell lines were analyzed by flow cytometry to assess GFP intensity per cell. **D.** GFP-RAB13 protein levels of the indicated cell lines were assessed by quantitative Western and normalized to α-Tubulin levels. n=7. n.s.: not significant.

**Figure S5:**
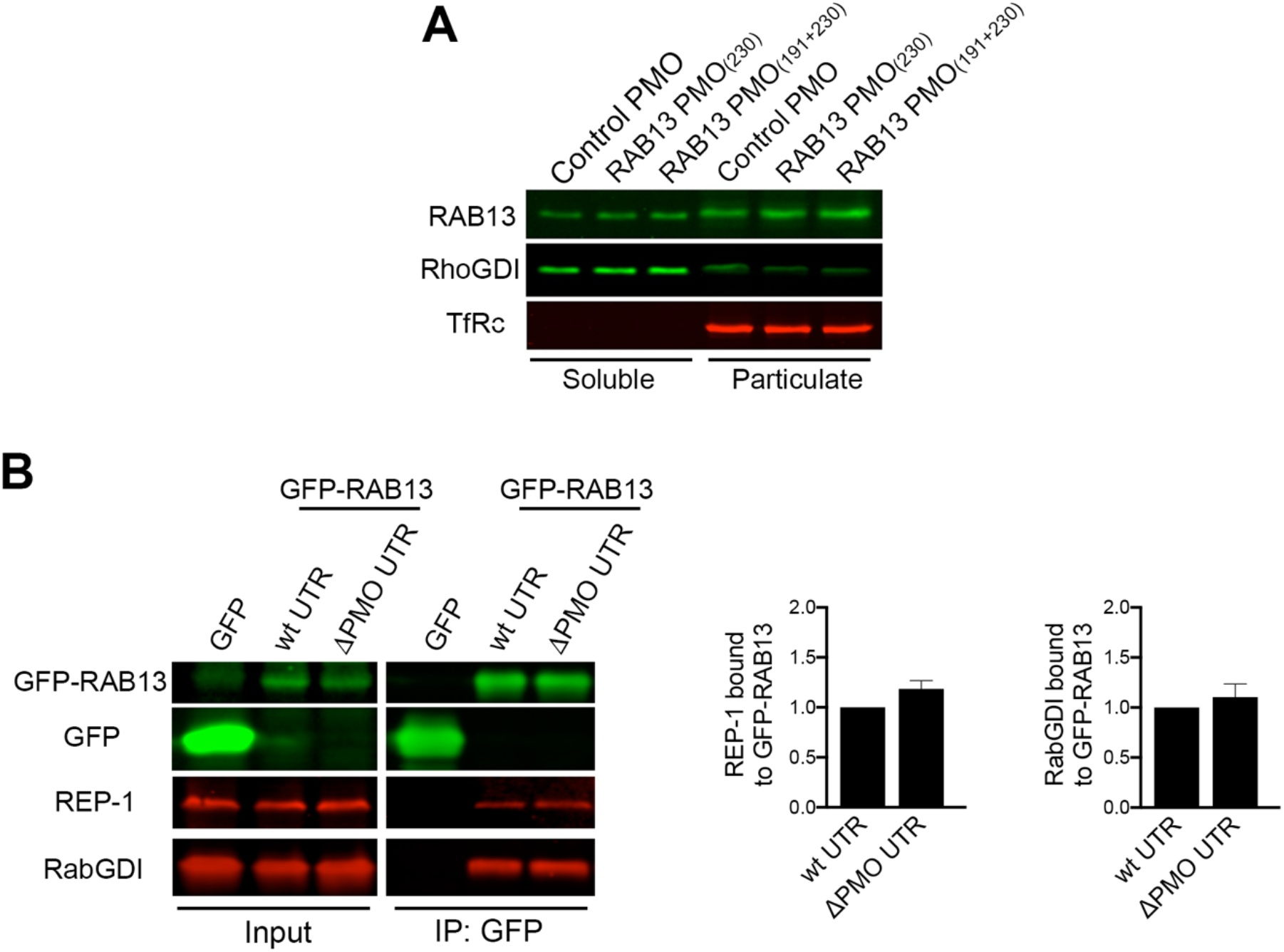
*RAB13* RNA mislocalization does not affect RAB13 binding to membranes or association with REP-1 or RabGDI. **A.** Cells treated with the indicated PMOs were fractionated into soluble and particulate fractions and the indicated proteins were detected by Western blot. RhoGDI and TfRc serve as soluble and particulate markers, respectively. **B.** Lysates from the indicated GFP or GFP-RAB13 expressing cell lines were immunoprecipitated with anti-GFP antibodies and blotted to detect the indicated proteins. Relative REP-1 and RabGDI binding is quantified in the graphs from n=3 (REP-1) and n=5 (RabGDI) independent experiments. No significant differences were detected.

**Figure S6:**
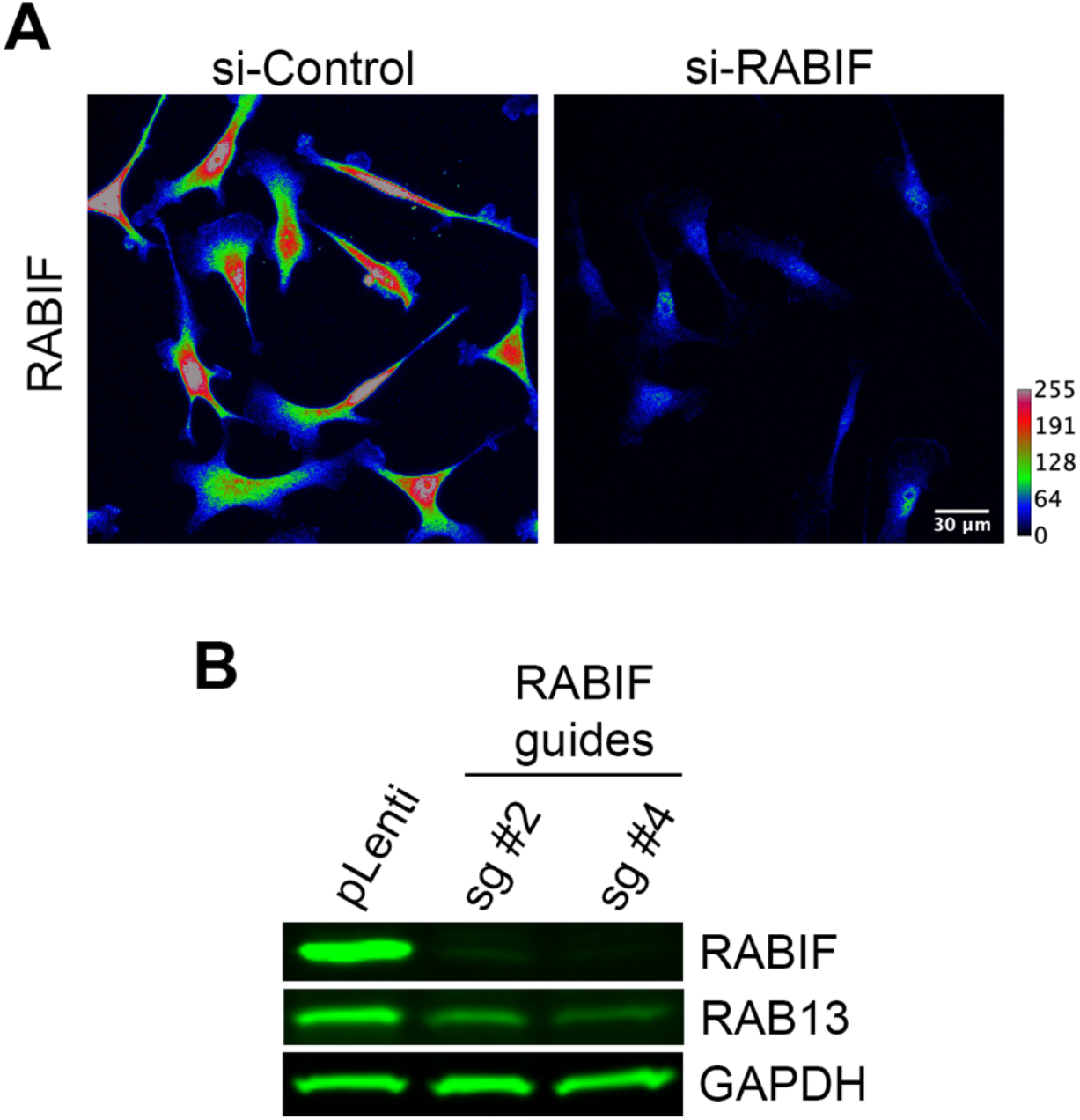
RABIF distribution and effect on RAB13 expression. **A.** Representative immunofluorescence images of RABIF protein in cells transfected with the indicated siRNAs. Reduction of intensity in knockdown cells confirms the specificity of the signal. Calibration bar shows intensity values. Note that RABIF exhibits a mostly perinuclear enrichment. **B.** RAB13 expression in cells with CRISPR knockdown of RABIF using the indicated sgRNAs (see also Figure 7C).

**Figure S7:**
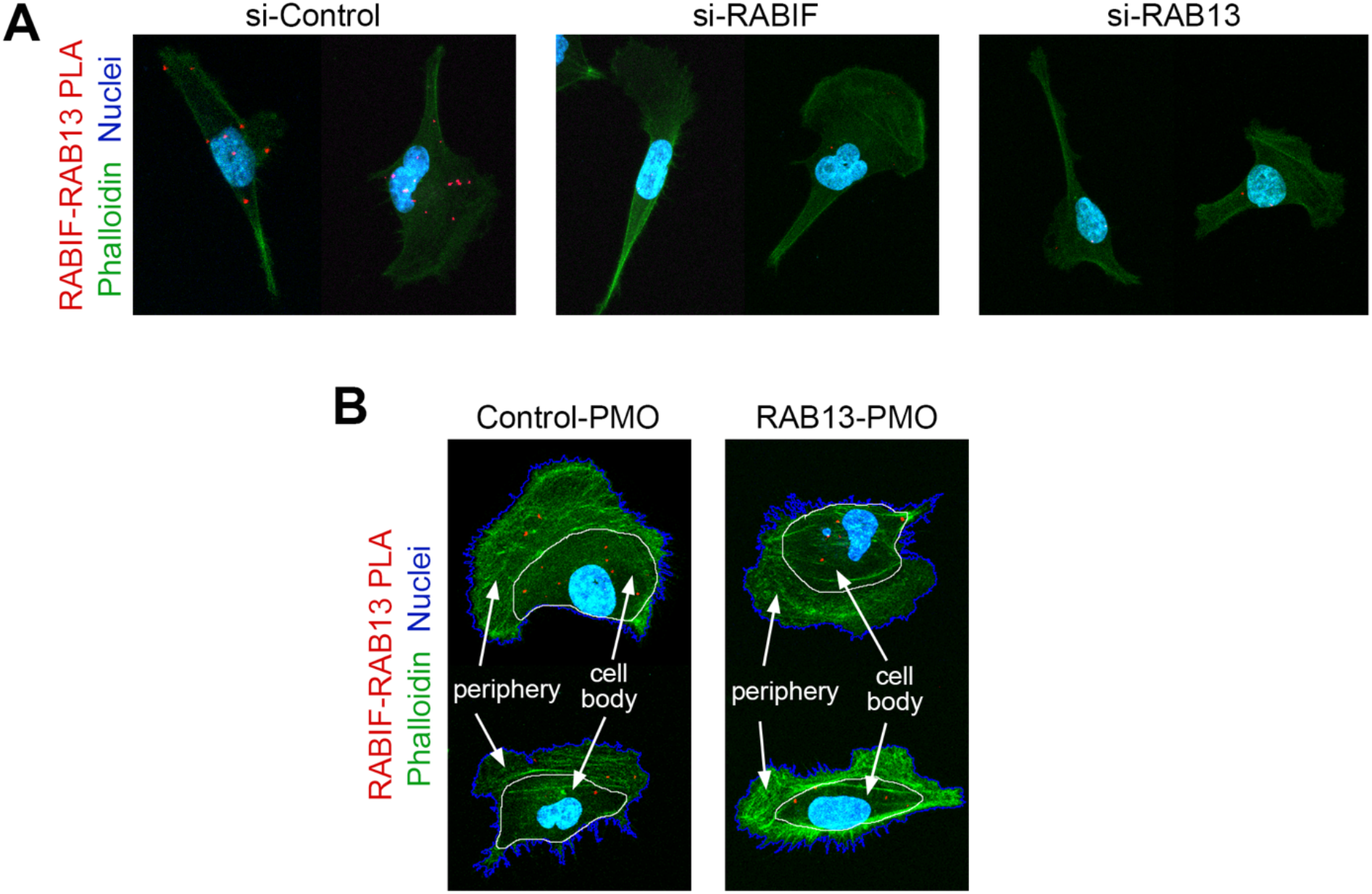
Detection of RABIF-RAB13 interaction through PLA. **A.** Representative RABIF-RAB13 PLA images from cells transfected with the indicated siRNAs (quantified in Figure 7D). **B.** Examples of sectioning of cell images into ‘peripheral’ and ‘cell body’ regions used for quantification of PLA signals in Figures 7E, F.

## SUPPLEMENTARY MOVIE LEGENDS

**Movie S1:** Representative time lapse imaging of MDA-MB-231 cells expressing Cherry-NLS to mark nuclei. Cells plated on collagen IV-coated coverglass were imaged over time and nuclei were tracked. Tracks of individual cells are overlaid in different colors. Time stamp in Hours:Minutes.

**Movie S2:** Left panel: Time lapse imaging of Lifeact-GFP-expressing cell plated on collagen IV-coated coverglass and imaged every minute over 1 hr. Right panel: Corresponding edge velocity map, with negative values indicating retraction and positive values indicating extension.

## SUPPLEMENTARY TALBLES

**Table S1: List of proteins interacting with GFP-RAB13(wt) or GFP-RAB13(T22N).** Table lists proteins identified by mass spectrometry analysis in two independent replicate experiments,

**Table S2: List of PMO oligonucleotides used in the study.**

